# The V-ATPase/ATG16L1 axis drives membrane remodeling during epithelial morphogenesis

**DOI:** 10.1101/2025.04.28.650675

**Authors:** Gabriel Baonza, Tatiana Alfonso-Pérez, Gonzalo Herranz, Carlos Quintana-Quintana, Carmen Gordillo-Vázquez, Yara El Mazjoub, L.M. Escudero, David G. Míguez, Elisa Martí, Nuria Martínez-Martín, Fernando Martín-Belmonte

**Affiliations:** Program of Tissue and Organ Homeostasis, Centro de Biología Molecular Severo Ochoa, Consejo Superior de Investigaciones Científicas-Universidad Autónoma de Madrid, Madrid 28049, Spain; Intestinal Morphogenesis and Homeostasis Group, Area 3-Cancer, Instituto Ramón y Cajal de Investigación Sanitaria (IRYCIS), Madrid 28034, Spain; Instituto de Biomedicina de Sevilla (IBiS), Hospital Universitario Virgen del Rocío/CSIC/Universidad de Sevilla. Departamento de Biología Celular, Facultad de Biología, Universidad de Sevilla, Seville 41013, Spain; Department of Cells and Tissues, Instituto de Biología Molecular de Barcelona, Parc Científic de Barcelona, Baldiri i Reixac 20, Barcelona 08028, Spain; Departmento de Física de la Materia Condensada, Instituto de Física de la Materia Condensada (IFIMAC), Facultad de Ciencias, Universidad Autónoma de Madrid, Madrid 28049, Spain

**Author notes:** These authors contributed equally (Gabriel Baonza, Tatiana Alfonso-Pérez).

**Keywords:** Epithelial Tubulogenesis, Autophagy, CASM, ATG16L1, V-ATPase, Human organoids, Tube micropatterns, Lumen resolution, Remodeling membranes, Endocytic trafficking

## Abstract

Epithelial tubulogenesis shapes internal organs by transforming flat epithelial sheets or unpolarized cords into hollow tubes with central lumens. A key example is the formation of the posterior neural tube during secondary neurulation, which requires precise morphogenetic events for *de novo* lumen formation. Although several studies have highlighted the role of autophagy in specific morphogenetic events, its involvement in epithelial organ development remains unclear. Autophagy operates via canonical and noncanonical pathways. Canonical autophagy is catabolic, requiring double-membrane autophagosomes and the full ATG protein set. Noncanonical autophagy, including the V-ATPase/ATG16L1-dependent Conjugation of ATG8 in Single Membranes (CASM), has both degradative and non-degradative roles and regulates different membrane trafficking processes. Using human neural tube organoids, spheroids, and tube micropatterns deficient in CASM or canonical autophagy, we show that CASM plays a pivotal role in epithelial tube morphogenesis. Specifically, the V-ATPase/ATG16L1 axis is essential for *de novo* lumen formation by regulating membrane junction remodeling and Rab11-dependent recycling pathways. These findings reveal distinct contributions of autophagy pathways in epithelial development, with potential implications for diseases linked to autophagy dysfunction.

## INTRODUCTION

The epithelium is a cellular tissue that covers all the exposed surfaces of the body. In a broad sense, it is the boundary that delimits the “inside” from the “outside”. While some epithelia cover the outer surface of the organism, most of it is organized internally in spaces or lumens forming tubular organs, where they serve to isolate and to perform fundamental functions such as excretion, gas exchange or digestion. Thus, in a prototypical epithelium, the apical surface faces the lumen, the lateral surface interacts with neighboring cells and the basal surface interacts with the extracellular matrix (ECM)^1–4^.

Strategies for *de novo* tube formation can be classified based on two distinct lumen formation mechanisms: (1) cavitation, where inner cells undergo apoptosis to create a luminal space, and (2) hollowing, where lumen formation involves directional vesicle trafficking and membrane separation without cell death^5^. The latter mechanism represents the intrinsic process by which many three dimensional (3D) cell models, such as Madin-Darby Canine Kidney (MDCK) spheroids, organoids and embryoids establish and maintain their 3D architecture^6–9^. In these models, cells inherently develop apical-basal polarity and undergo stereotypical morphogenesis, transitioning from single cells to a polarized monolayer surrounding a central lumen. Upon 3D plating, cells initially form doublets with inverted polarity, where apical proteins like Podocalyxin (Podxl) localize to the ECM-facing surface. Integrin-mediated mechanosensing triggers the endocytosis and transcytosis of Podxl and other apical proteins to the apical membrane initiation site (AMIS), which subsequently remodels into the nascent lumen. This process reworks basolateral domains into a specialized apical delivery zone, the pre-apical patch (PAP), facilitating lumen expansion and maturation^7,10^. Compellingly, despite the diversity of mechanisms involved, the fundamental principle of cell polarity remains central to successful lumenogenesis. In particular, polarized membrane transport has emerged as a key regulatory pathway, crucial not only for establishing individual epithelial cell polarity but also for coordinating epithelial polarization during apical lumen formation and tissue morphogenesis^11–13^.

It has been described that during lumenogenesis, cells undergo a phase of enhanced metabolism and architectural changes, where they modify their organelle and protein content to adapt and respond to altering conditions. Since this phase is very brief, proteasome-mediated degradation of proteins is probably not enough to compete with the occurring architectural changes. In this sense, autophagy, which is faster and more versatile, could contribute to this process^14,15^.

Autophagy is an evolutionary, highly conserved bulk degradation mechanism used by all eukaryotic cells to remove or recycle intracellular components. Commonly referred to as macroautophagy, this catabolic process is typically recognized for alleviating cellular stress triggered by low cellular nutrient levels, the accumulation of misfolded or long-lived proteins, damaged organelles, aggregates or pathogens. During macroautophagy, a cohort of different ATG proteins orchestrates the initiation, expansion, and maturation of double-membrane vesicles known as autophagosomes, which encapsulate target cargo for subsequent lysosomal degradation. A defining hallmark of this process is the covalent lipidation of LC3 (conversion of LC3-I to LC3-II) and its insertion into the expanding phagophore, a step mediated by the ATG12-ATG5-ATG16L1 complex^15^. Intriguingly, a subset of ATG proteins operates in parallel to mediate noncanonical autophagy, a pathway that specifically targets single-membrane endolysosomal compartments^16–20^. Examples of this noncanonical autophagy include pathogen clearance, membrane repair, and unconventional protein secretion through secretory autophagy^21–23^. While protein transport is a well-established regulator of lumen formation and function, and the interplay between endolysosomal and autophagosomal pathways has been extensively studied in mammalian cells, the precise role of autophagy in lumenogenesis remains largely unexplored.

In the present study, we investigate the role of autophagy in key events of epithelial lumen formation, highlighting its functional diversity in morphogenetic processes. We propose that noncanonical autophagy, is activated during epithelial morphogenesis to facilitate membrane remodeling, thereby contributing to the establishment of a single, functional lumen. Additionally, we demonstrate that canonical autophagy regulates hydrostatic pressure by modulating ion channel activity, which is essential for lumen expansion. Finally, we delineate distinct mechanisms through which both canonical and noncanonical autophagy contribute to lumenogenesis, underscoring their coordinated roles in shaping epithelial architecture.

## RESULTS

### Autophagy is essential for lumen resolution during human neural tube secondary neurulation

Secondary neurulation is a fundamental morphogenetic process responsible for forming the caudal region of the neural tube in vertebrates. Unlike primary neurulation, which involves folding and fusion of the neural plate, secondary neurulation proceeds through a mesenchymal-to-epithelial transition (MET), followed by cavitation within a solid medullary cord^24–26^. During this process, multiple small lumen foci emerge *de novo*, which subsequently coalesce to form a single central lumen. The sequential exposure of neural progenitor cells to signaling pathways—including TGFβ/BMP, WNT, FGF, and retinoic acid (RA)—regulates spatiotemporal gene expression, promoting lineage specification and cellular organization^7^. Notably, these key features of secondary neurulation can be faithfully recapitulated in vitro using human neural tube organoids (huNTOrgs)^7^, which serve as an accessible and tractable model to study posterior neural tube morphogenesis (Figure 1A).

**FIGURE 1.**
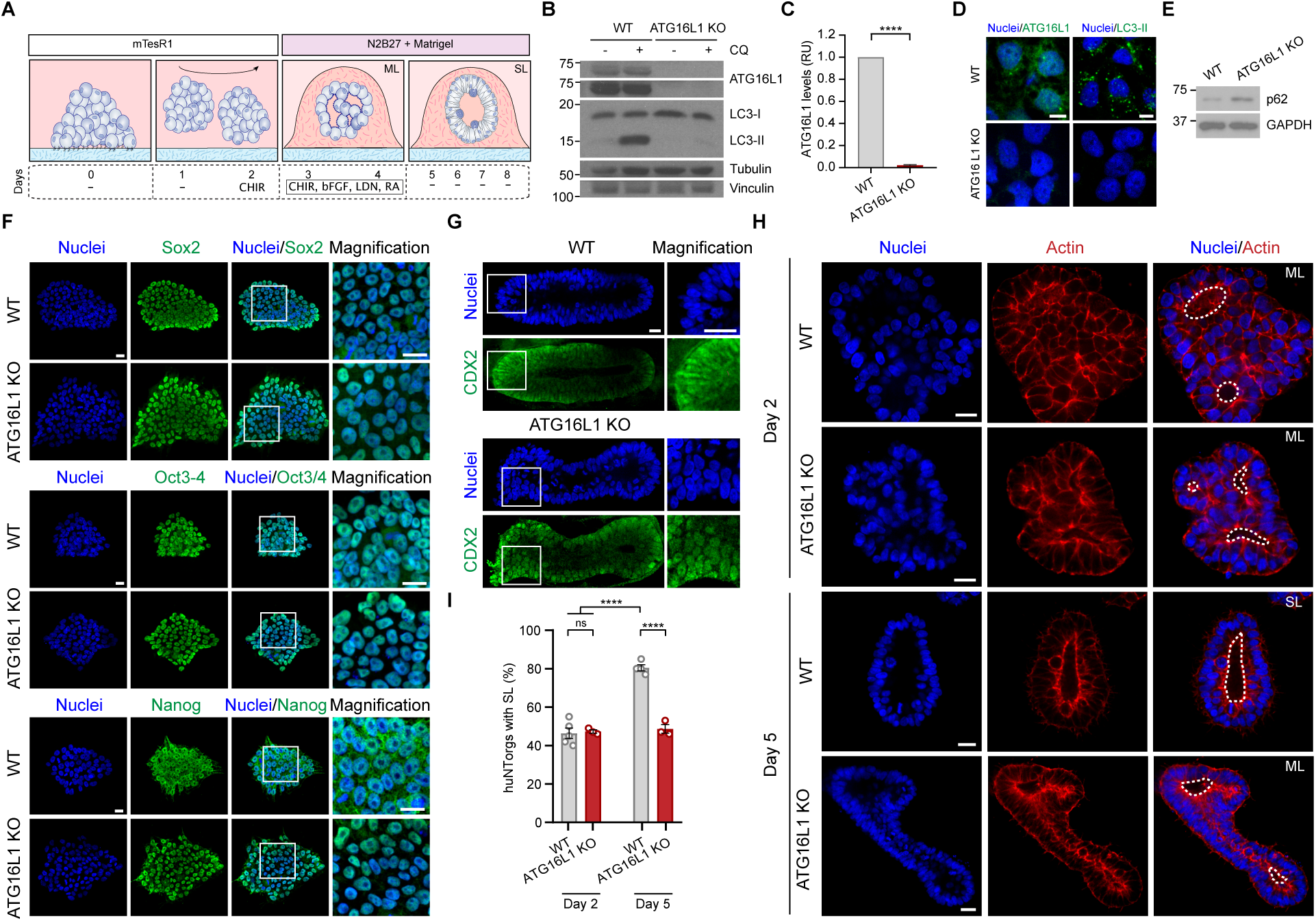
ATG16L1 KO impairs lumen resolution during human neural tube secondary neurulation. **(A)** Schematic representation of the conditions used to model lumen resolution during human neural tube secondary neurulation. ML=multi lumen. SL=single lumen. **(B)** Absence of ATG16L1 impairs LC3 lipidation in ATG16L1 knockout (KO) H1 hESCs, assessed by western blot (WB). +/-=with/without. Chloroquine (CQ). **(C)** Quantification of ATG16L1 protein levels in WT and ATG16L1 KO H1 hESCs. Relative units (RU). Data are expressed as the mean ± SEM from 3 independent experiments. Unpaired two-tailed t-test was conducted. **(D)** Immunofluorescence (IF) shows absence of ATG16L1 and LC3-II in ATG16L1 KO H1 hESCs. Scale bar=10 μm. **(E)** Increased p62 protein level in ATG16L1 KO H1 hESCs, analyzed by WB. **(F)** Expression of pluripotency markers Sox2, Oct3/4, and Nanog in WT and ATG16L1 KO H1 hESCs. **(G)** IF shows expression of CDX2 in posterior human neural tube organoids (huNTorgs) derived from WT and ATG16L1 KO H1 hESCs at day 5 of development. **(H)** ATG16L1 KO H1 huNTorgs fail to resolve ML at day 5 of development. Cells were stained for actin (red) and nuclei (blue). Dotted lines indicate the lumens. **(I)** Percentage (%) of SL quantified from IF images shown in (B). Data are expressed as the mean ± SEM from at least 3 independent experiments. (N=1022 organoids). One-way ANOVA test was performed. In panels D, F, G, and H nuclei are stained with DAPI. In panels F-H, Scale bar=20 μm. *P* values=\*\*\*\**p*<0,0001; ns=non-significant.

Given the pivotal role of autophagy in various morphogenetic contexts, we sought to investigate its specific contribution to secondary neurulation. Autophagy encompasses two distinct pathways: canonical and noncanonical, both of which converge on ATG16L1, a core component of the autophagic machinery^20,27^. To dissect the role of autophagy in neural tube formation, we first established a complete loss-of-function model by generating homozygous ATG16L1 knockout (KO) human embryonic stem cells (hESCs) via CRISPR/Cas9-mediated genome editing (Figure 1B). Following genome editing, we confirmed the complete absence of ATG16L1 protein expression by western blotting (Figure 1B-1C) and observed the expected loss of LC3 lipidation (conversion of LC3-I to LC3-II) after chloroquine (CQ) treatment (Figure 1B). CQ, a lysosomal inhibitor, blocks the fusion of autophagosomes with lysosomes, thereby preventing the degradation of autophagic substrates^28^. Immunofluorescence analysis further validated the absence of ATG16L1 protein levels and LC3 puncta (Figure 1D). Consistent with a block in autophagic flux, we detected increased levels of the autophagy substrate p62/SQSTM1^29^ (Figure 1E). Together, these results confirmed the successful establishment of a model with complete impairment of autophagy.

Before proceeding with neural differentiation, we examined the pluripotent state of the ATG16L1 KO hESCs. Morphologically, KO colonies displayed typical features of undifferentiated hESCs, including compact colony architecture, high nucleus-to-cytoplasm ratios, and prominent nucleoli^30^ (Figure S1A). Immunofluorescence staining demonstrated nuclear localization of key pluripotency markers Sox2, Oct3/4, and Nanog^31^, with expression levels comparable to wild-type (WT) controls (Figure 1F). Flow cytometric analysis further confirmed these findings by assessing both transcription factors and surface markers, including SSEA4 and SSEA1^32^, the latter indicative of early differentiation (Figure S1B). Additionally, genome-wide analysis of copy number variations revealed no abnormalities, ruling out potential clonal artifacts (Figure S1C). These results collectively established that ATG16L1 KO hESCs retained pluripotency and genomic stability.

Having validated the ATG16L1 KO cell line, we next sought to investigate the impact of autophagy loss on huNTOrgs development. Both WT and ATG16L1 KO hESCs were guided through neural induction protocols to generate huNTOrgs with posterior identity (Figure 1A). In this in vitro model of secondary neurulation, cell aggregates are first cultured under agitation. Transference to Matrigel promotes MET and small lumen formation, that later coalesce in a single lumen, when cultured in neural induction medium^7^. As expected, WT huNTOrgs exhibited robust posterior identity, as demonstrated by the expression of the caudal marker CDX2^33^, and successfully underwent lumen resolution, forming a single central lumen within five days of differentiation (Figures 1G). Strikingly, ATG16L1 KO huNTOrgs retained CDX2 expression (Figure 1G) but consistently failed to resolve multiple lumens into a single central cavity (Figures 1H-1I, S1D and S1F). Quantitative image analysis revealed that, in contrast to WT organoids, nearly 50% of ATG16L1 KO huNTOrgs retained a persistent multilumen phenotype after five days of culture (Figures 1H-1I). These findings suggest that autophagy plays a critical role in facilitating the resolution of multiple lumens during secondary neurulation.

To further explore whether this requirement for autophagy is specific for lumen resolution or rather general for organoid morphogenesis, we analyzed organoids derived from ATG16L1 KO mouse ESCs. Unlike huNTOrgs, mouse ESC-derived organoids (blastuloids) do not form multiple lumen intermediates but instead generate a single lumen through an alternative mechanism that bypasses the need for lumen coalescence^34,35^ (Figure S2A). Consistent with this difference, ATG16L1 KO mouse organoids did not exhibit defects in lumen formation (Figures S2A-S2E). These observations suggest that autophagy-dependent lumen resolution may represent a specific mechanism during posterior neural tube development.

In summary, these results highlight an essential role for autophagy in the resolution of multiple lumens during human secondary neurulation. The failure of this process in ATG16L1 KO huNTOrgs implicates autophagy dysfunction in the pathogenesis of neural tube defects and underscores the need for further experiments to elucidate the molecular mechanisms underpinning this process.

### Canonical Autophagy Is Dispensable for Lumen Resolution During Epithelial Morphogenesis

Our previous findings demonstrate that the complete loss of ATG16L1 impairs lumen resolution during human secondary neurulation. Yet, given the fact that ATG16L1 functions in both canonical and noncanonical autophagy pathways^20,27^, we sought to further dissect and uncover which of these mechanisms underlies the observed phenotype. To address this question and validate our observations in an alternative epithelial model, we turned to the 3D MDCK organotypic culture system. This well-established model faithfully recapitulates key aspects of epithelial lumen formation and tissue architecture, while offering advantages in terms of manipulation, reproducibility, and visualization^1^.

We first generated multiple homozygous ATG16L1 KO MDCK cell lines using a CRISPR/Cas9 genome editing approach (Figure S3A). These clones were characterized at various stages of 3D cyst development, spanning from 24 to 72 hours of culture. Western blot analysis confirmed a complete loss of ATG16L1 protein in KO clones (Figure 2A-2B). Consistent with a disruption in autophagy, we observed a total abrogation of LC3 lipidation (conversion of LC3-I to LC3-II), and an accumulation of the selective autophagy receptor p62/SQSTM1 (Figures 2A-2B). Additionally, we detected reduced expression of WD repeat domain phosphoinositide-interacting protein 2 (WIPI2) (Figures S3A-S3B), a key component involved in the early stages of autophagosome formation^36^. To further confirm these findings at the cellular level, we performed confocal microscopy on both WT and ATG16L1 KO MDCK cysts at 72 hours of development. As was foreseeable, WT cysts exhibited normal ATG16L1 expression localized to the appropriate cellular compartments, whereas ATG16L1 KO cysts displayed a complete absence of the protein (Figure 2C).

**FIGURE 2.**
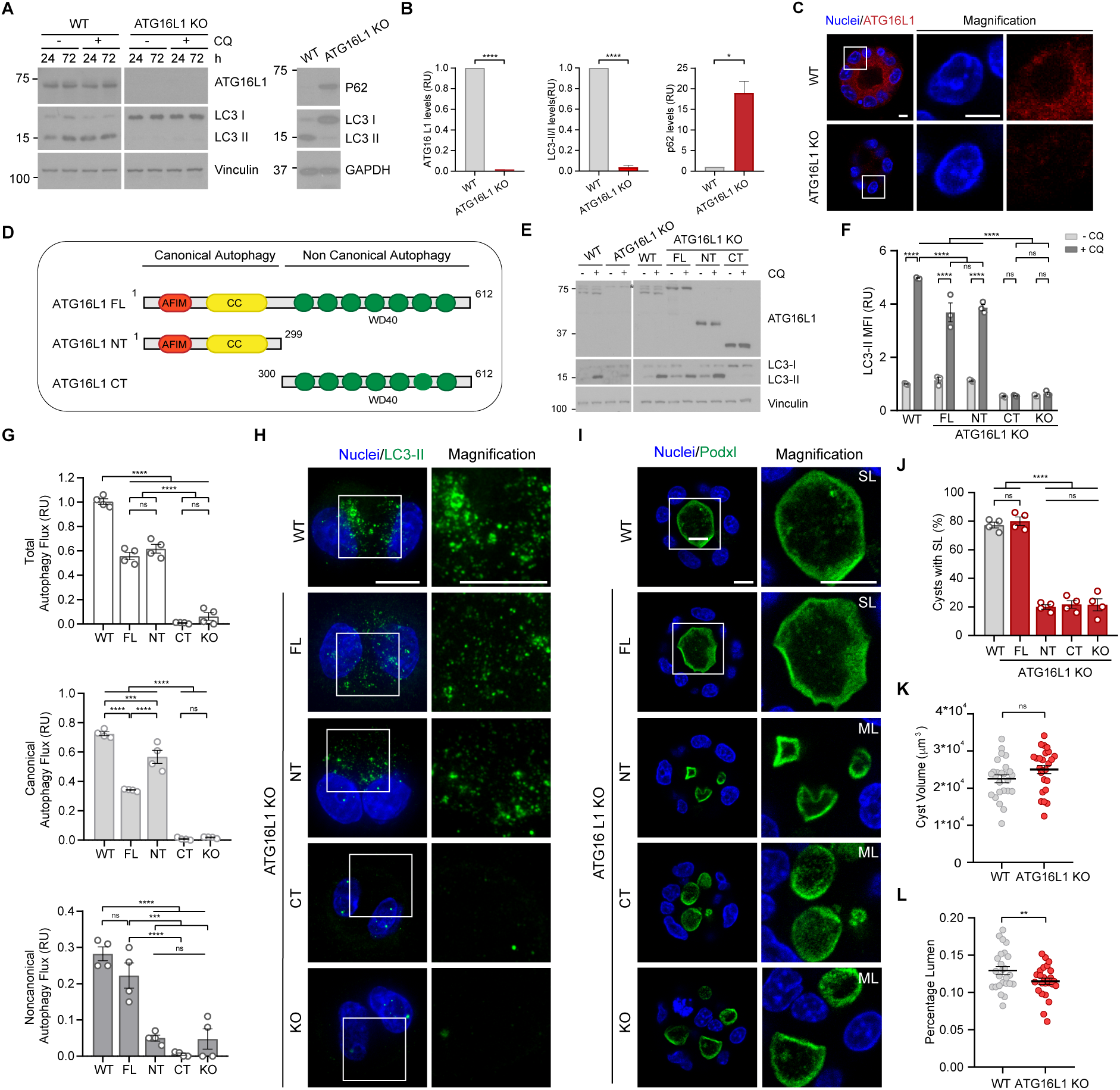
Noncanonical autophagy is critical to resolve multi lumens into a single cavity during epithelial morphogenesis. **(A)** Absence of ATG16L1 impairs LC3 lipidation and increases p62 in 24-72 hours (h) cysts derived from ATG16L1 KO cells, assessed by WB. With or without (+/-). **(B)** Quantification of ATG16L1, LC3-II/LC3I and p62 in ATG16L1 KO compare to 72h WT cysts. Relative units (RU). Data are expressed as the mean ± SEM from 3 independent experiments. Unpaired two-tailed t-test was conducted. **(C)** The absence of ATG16L1 in 72h cysts derived from ATG16L1 KO cells. Scale bar=5 μm. **(D)** Schematic representation of ATG16L1 constructs: full-length (FL), amino-terminal (NT), and carboxyl terminal (CT). **(E)** ATG16L1 expression and LC3 lipidation in WT and ATG16L1 KO cells stably expressing FL, NT, or CT constructs, assessed by WB. **(F)** LC3-II mean fluorescence intensity (MFI) quantified by flow cytometry in 24h MDCK cysts from WT, ATG16L1 KO, FL-ATG16L1 KO, NT-ATG16 KO, and CT-ATG16L1 KO cells lines. Relative units (RU). Data are expressed as the mean ± SEM from 3 independent experiments. Two-way ANOVA test was performed. **(G)** Autophagy Flux represented by LC3II MFI values, quantified by flow cytometry in 24h cysts derived from WT, FL-ATG16L1 KO, NT-ATG16 KO, CT-ATG16L1 KO, and ATG16L1 KO cell lines. Data are expressed as the mean ± SEM from 4 independent experiments. One-way ANOVA was performed. **(H)** 24h cysts show LC3 lipidation in WT, FL-ATG16L1 KO, and NT-ATG16L1 KO, whereas no LC3 lipidation is observed in ATG16L1 KO or CT-ATG16L1 KO cysts in the presence of CQ. Scale bar=10 μm. **(I)** 72h cysts shows that ATG16L1 KO, NT-ATG16L1 KO, and CT-ATG16L1 KO fail to resolve multi lumens (ML). SL=single lumen. Podocalyxin (Podxl). Scale bar=10 μm. **(J)** Percentage (%) of SL quantified from IF images shown in (I). Data are expressed as the mean ± SEM from 4 independent experiments (N=2000 cysts). One-way ANOVA test was performed. **(K)** WT and ATG16L1 KO 72h cysts with single lumen exhibit similar cyst volume (µm³). **(L)** ATG16L1 KO 72h cysts with single lumen exhibit less percentage lumen. In panels K-L data are expressed as the mean ± SEM from 3 independent experiments (N=50 cysts) and quantified using CartoCell. Unpaired two-tailed t-test was conducted. In panels C and H-I, nuclei are stained with DAPI (blue). *P* values=\*\*\*\**p*<0.0001; \*\*\**p*<0.001; \*\**p*<0.01; ns=non-significant.

Taken together, these results confirm that ATG16L1 KO MDCK cysts exhibit a profound and complete impairment of the autophagy pathway, mirroring the observations made in our ATG16L1 KO huNTOrgs. This provides a robust platform to further interrogate the specific role of autophagy—particularly canonical versus noncanonical pathways—in regulating epithelial lumen resolution.

ATG16L1 plays a dual role in autophagy, orchestrating both canonical and noncanonical pathways through distinct domains (Figure 2D). Structurally, ATG16L1 is organized into three regions: an N-terminal (NT) segment bearing the ATG5-interacting motif (AFIM), a central coiled-coil (CC) domain, and a C-terminal WD40 region of seven β-propeller repeats. The NT and CC segments together form the N-terminal domain, while the WD40 repeats constitute the C-terminal domain. The NT region is essential for LC3 lipidation in both pathways by recruiting the ATG12-ATG5 to the complex^37,38^. In canonical autophagy, this process requires interaction with WIPI2 and facilitates autophagosome formation on double membranes. Conversely, the CT WD40 domain is crucial for noncanonical autophagy, enabling LC3 lipidation on single membranes independently of WIPI2 by interacting with V-ATPase^39,40^. This structural division highlights a context-dependent coordination between ATG16L1 domains in regulating autophagic processes.

Accordingly, we aimed to dissect the dual functionality of ATG16L1 by distinguishing the contributions of its NT and CT domains. To this end, we generated three stable MDCK cell lines, each expressing either the NT domain, the CT domain, or the full-length (FL) ATG16L1 protein, in the background of ATG16L1 KO MDCK cells (Figure 2D). We first evaluated LC3 lipidation in these cell lines, comparing WT, ATG16L1 KO, ATG16L1 KO + FL, ATG16L1 KO + NT, and ATG16L1 KO + CT cells. Western blot analysis showed robust LC3-II accumulation in WT, FL-, and NT-expressing ATG16L1 KO cells, while neither the ATG16L1 KO cells nor those expressing the CT domain exhibited LC3-II accumulation (Figure 2E). These findings underscore the critical role of the NT domain of ATG16L1 in mediating LC3 lipidation, consistent with its known function in canonical autophagy. To further quantify LC3 lipidation, we performed flow cytometry analysis of MDCK cysts after 24 hours of development. The results mirrored those from the western blot, confirming LC3-II accumulation in FL-and NT-expressing cells, albeit at levels noticeably lower than in WT cells (Figure 2F). These findings suggest that the NT domain of ATG1L1 is sufficient to mediate LC3 lipidation.

We then assessed the autophagic flux in FL-, Nt-, and Ct-expressing ATG16L1 KO cells by evaluating LC3-II accumulation following treatment with two widely used autophagy inhibitors, bafilomycin A1 (BAF) and CQ. These inhibitors block LC3-II degradation through distinct mechanisms, thereby enabling us to distinguish between canonical and noncanonical autophagy pathways^21^. Not surprisingly, both ATG16L1 KO and CT-ATG16L1 KO cells displayed a complete absence of autophagic flux under all treatment conditions, underlining their inability to lipidate LC3 (Figures 2G and S3C). In contrast, while FL-ATG16L1 KO cells showed a slight reduction in total autophagic activity compared to control cells, the relative balance between canonical and noncanonical was largely restored (Figure 2G). Notably, NT-ATG16L1 KO cells presented similar autophagic flux when compared to FL-ATG16L1 KO or control cells, which is provided mostly by an increased canonical autophagy, complementing the significant reduction of noncanonical autophagy (Figure 2G). In line with these findings, immunofluorescence analysis of MDCK cysts generated from all cell lines yielded results consistent with those obtained by flow cytometry. Specifically, LC3 lipidation was observed after 24 and 72 hours of cyst development and CQ treatment in controls, and the NT- or FL-versions of ATG16L1 (Figures 2H and S3D).

The outcomes obtained through the preceding approaches are congruent with previous reports^41^ and align with the functional architecture of ATG16L1. Only the FL- and NT-constructs retain the ATG5-binding domain, essential for ATG12-ATG5-ATG16L1 complex formation and subsequent LC3 lipidation. Accordingly, both canonical and noncanonical autophagy were rescued in FL-expressing ATG16L1 KO cells, whereas only canonical autophagy was restored in ATG16L1 KO cells expressing the N-terminal fragment of ATG16L1.

After establishing models that differentiate between cells exhibiting both canonical and noncanonical autophagy (WT and FL-expressing cells), canonical autophagy alone (NT-expressing cells), or the complete absence of autophagy (CT-expressing and KO cells), we addressed whether autophagy is required for lumenogenesis in MDCK cysts. To investigate this, cysts from all cell lines were cultured for 72 hours and stained for Podxl to visualize and quantify lumen formation. As expected, depletion of ATG16L1 profoundly impaired single-lumen formation, resulting in the frequent emergence of multiple ectopic lumens (Figures 2I-J and S3E). This phenotype closely resembled the impaired lumen resolution previously observed in posterior huNTOrgs derived from ATG16L1 KO hESCs (Figures 1H–1I). Interestingly, even in the subset of ATG16L1 KO cysts that successfully formed a single lumen, lumen circularity and expansion were significantly compromised (Figures S3F–S3G). High-content segmentation analysis using CartoCell^42^, a novel image quantification framework, revealed that although these cysts contained the same number of cells as their WT counterparts, the epithelial cell height was consistently higher in ATG16L1 KO cysts (Figures S3H-S3I). This observation is consistent with the reduced lumen expansion phenotype in ATG16L1 KO cysts capable of forming a single lumen (Figures 2K–2L). Strikingly, only FL-expressing ATG16L1 KO cysts—where both canonical and noncanonical were restored—successfully rescued the single-lumen phenotype. In contrast, NT-expressing ATG16L1 KO cysts, in which only canonical was present, failed to restore proper lumen formation and exhibited a persistent multilumen phenotype (Figures 2I–2J).

These observations demonstrate a distinct functional divergence between full-length ATG16L1 and its isolated N-terminal domain in autophagy-mediated morphogenesis. While complete autophagy impairment severely disrupts lumen resolution, canonical autophagy alone appears dispensable for this process. Instead, noncanonical autophagy mediated by the C-terminal WD40 domain of ATG16L1 plays a critical role in ensuring proper lumen morphogenesis during epithelial tubulogenesis.

### Noncanonical Autophagy Is Essential for Lumen Resolution, While Canonical Autophagy Contributes to Lumen Expansion During Epithelial Morphogenesis

The inability of canonical autophagy alone to rescue the multilumen phenotype—while restoration of a single, central lumen requires the concurrent presence of both canonical and noncanonical autophagy pathways—strongly suggests that noncanonical autophagy plays a paramount function in epithelial lumen formation. Recent studies have identified the vacuolar ATPase (V-ATPase) proton pump as a key regulator of noncanonical autophagy, specifically CASM. In particular, its V1H subunit directly interacts with ATG16L1 within the ATG12–ATG5–ATG16L1 complex, but only when the V-ATPase is fully assembled following organelle neutralization (Figure S4A). This interaction is essential for targeting LC3 to single-membrane compartments—a defining feature of CASM. While many V-ATPase subunits are essential for viability, ATP6V1H-deficient cells remain viable but are unable to recruit LC3 to these compartments, indicating a specialized role for this subunit in noncanonical autophagy^39,43,44^. On the other hand, depletion of WIPI2 disrupts canonical autophagy while leaving noncanonical autophagy intact. This is supported by extensive literature showing that noncanonical autophagy bypasses certain upstream regulators, such as WIPI2, though it still relies on the core ubiquitin-like conjugation system for LC3 lipidation^21,45^.

Based on these distinctions, we generated two complementary MDCK KO models using CRISPR/Cas9: one lacking ATP6V1H (disrupting noncanonical autophagy), and the other lacking WIPI2 (disrupting canonical autophagy) (Figures S4B-S4C). Interestingly, although both ATP6V1H and WIPI2 KO cells maintained LC3 lipidation during cyst development—as demonstrated by western blot and flow cytometry (Figures 3A–3B and S4C)—their autophagic flux profiles diverged significantly (Figure 3C). Specifically, ATP6V1H-deficient cysts maintained canonical autophagy flux at levels similar to WT cells, while WIPI2-deficient cysts relied solely on noncanonical autophagy, exhibiting more than twice the activity of WT controls. This enhanced noncanonical autophagy in WIPI2 KO cysts was accompanied by prominent co-localization of V-ATPase and ATG16L1 at shared compartments after 24 hours—a hallmark of CASM activity^43,46^ (Figure 3D). Further supporting this, p62 levels were elevated in both ATG16L1 and WIPI2 KO cysts, indicating impaired canonical autophagy. In contrast, ATP6V1H and NT-ATG16L1 KO cysts displayed lower p62 accumulation, consistent with preserved canonical function (Figures 3E–3F). Moreover, WIPI2/ATP6V1H double KO cysts exhibited a complete absence of LC3 lipidation and autophagic activity, validating the pathway-specific roles of these two factors (Figure S4D).

**FIGURE 3.**
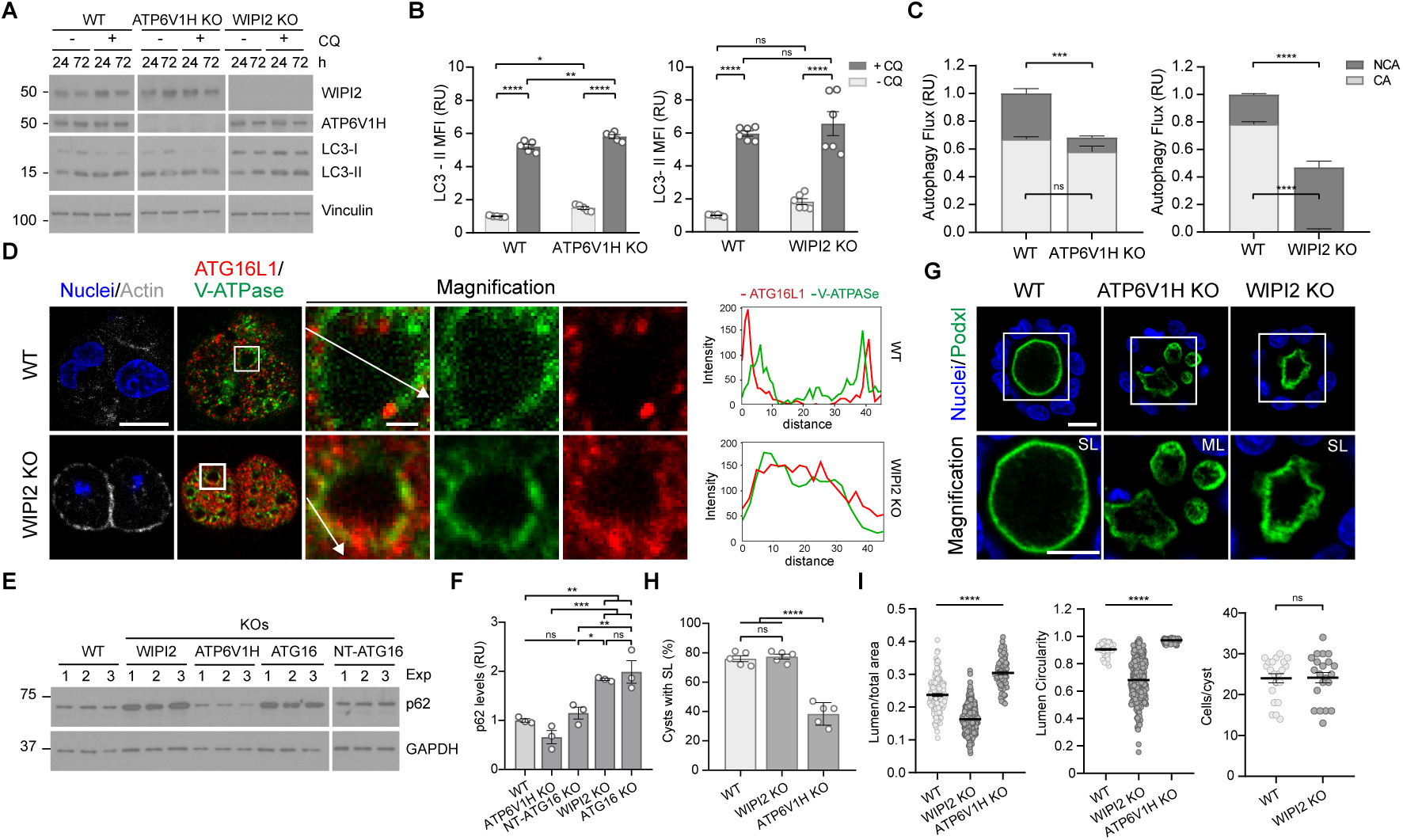
Disruption of CASM autophagy (ATP6V1H KO) impairs lumen resolution, while loss of canonical autophagy (WIPI2 KO) affects lumen expansion. **(A)** Absence of ATP6V1H or WIPI2 does not impair LC3 lipidation in 24-72h cysts, assessed by WB with or without (-/+) Chloroquine (CQ). **(B)** LC3-II MFI quantified by flow cytometry in 24h cysts from WT, ATP6V1H KO, and WIPI2 KOs cells lines. Data are expressed as the mean ± SEM from at least 5 independent experiments. Two-way ANOVA test was performed. **(C)** Loss of ATP6V1H disrupts noncanonical autophagy (NCA), loss of WIPI2 disrupts canonical autophagy (CA). Autophagy Flux represented by LC3II MFI values, quantified by FACs in 24h cysts derived from WT, ATP6V1H KO, and WIPI2 KO cells. Data are expressed as the mean ± SEM from 5 independent experiments. Unpaired two-tailed t-tests was conducted. **(D)** Co-localization of ATG16L1 and V-ATPase at vesicle structures in 24h cysts derived from WT and WIPI2 KO cells treated with CQ. Scale bars =10 μm and 1 μm (magnification) (in right panels). Fluorescence intensity profile is plotted along the white lines (right). **(E)** p62 protein levels increase in ATG16L1 KOs and WIPI2 KOs, but remain unchanged in WT, ATP6V1H KO, and NT-ATG16L1 KO at 72h cyst development, assessed by WB. **(F)** Quantification of p62 levels from WBs shown in panel (E). Data are expressed as the mean ± SEM from 3 independent experiments. One-way ANOVA test was performed. **(G)** 72h cysts shows that ATP6V1H KO cells fails to resolve multiple lumens (ML) and WIPI2 KO cells exhibits smaller lumen cavities. Podocalyxin (Podxl). Scale bar=10 μm. **(H)** Percentage (%) of SL quantified from IF images shown in (G). Data are expressed as the mean ± SEM from 5 independent experiments (N=1500 cysts). One-way ANOVA test was performed. **(I)** Quantitative analysis showing that WIPI2 KO 72h cysts exhibit smaller lumen area and reduced lumen circularity, without any change in cell number. Data are expressed as the mean ± SEM from 3 independent experiments. Kruskal-Wallis test was performed to analyze lumen and circularity data. Unpaired two-tailed t-test was performed to analyze cells/cyst. (N=525 cyst for lumen area, N=529 cysts for circularity, N=42 cysts for cells/cyst). In panels D and G, nuclei are stained with DAPI (blue). In panels B, C, and F, Relative units (RU). *P* values= \*\*\*\**p*<0.0001; \*\*\**p*<0.001; \*\**p*<0.01, and \**p*<0.05; ns=non-significant.

We then examined the functional impact of each pathway on epithelial lumen formation. Notably, ATP6V1H KO cysts failed to establish a single central lumen and instead developed multiple lumens, phenocopying the defect seen in ATG16L1 KO organoids (Figures 3G–3H). Conversely, WIPI2 KO cysts retained the ability to form single lumens, but exhibited significantly smaller luminal cavities and reduced circularity after 72 hours, without any change in total cell number (Figure 3G-3I and S4E–S4G).

Given the known role of Claudin-2 (Cldn2) in regulating epithelial permeability and lumen expansion, we examined its expression^47^. Claudin-2 levels were markedly reduced in WIPI2 and ATG16L1 KO cysts— both deficient in canonical autophagy—whereas ATP6V1H KO cysts preserved its expression (Figures S4H–S4I). Since previous studies have linked canonical autophagy to Claudin-2 regulation via p62-mediated degradation^48^, the reduction observed here may underlie the impaired lumen expansion seen in these cells.

Together, these findings reveal a division of labor between the autophagy pathways: noncanonical autophagy is essential for resolving multiple lumens into a single central cavity, while canonical autophagy primarily contributes to lumen expansion and epithelial structural refinement.

Noncanonical Autophagy is mediated by large, non-degradative, LC3-positive vesicles^16,17^. Having established the essential role of noncanonical autophagy in lumen resolution during epithelial morphogenesis, we next sought to characterize the identity of the vesicular structures associated with this unconventional pathway. To this end, we focused on immunofluorescence-based characterization of vesicles in WIPI2-deficient cysts, which exclusively exhibit noncanonical autophagy. These were compared to ATP6V1H KO and NT-ATG16L1 KO cysts, which exhibit only canonical autophagy, as well as to ATG16L1 KO cysts (autophagy-deficient) and WT controls (Figure 4A-4B). Vesicle size analysis revealed that WIPI2 KO cysts consistently exhibited large LC3-positive vesicles at two key developmental stages: 24 hours, during polarity establishment and AMIS formation, and 72 hours, during lumen resolution and expansion. In contrast, ATP6V1H and NT-ATG16L1 KOs predominantly displayed small LC3-positive vesicles, suggesting impaired vesicle maturation. WT cysts exhibited a heterogeneous population of vesicles, with both small and large LC3-positive structures present. Notably, ATG16L1 KO cysts showed a complete absence of LC3 signal, indicating a total loss of autophagic activity. Moreover, treatment with BAF, which inhibits V-ATPase and selectively abolishes noncanonical autophagy, led to the disappearance of large LC3-positive vesicles in both WIPI2 KO and WT cysts (Figure 4C). Together, these findings suggest that noncanonical autophagy is associated with large LC3-positive vesicles, while canonical autophagy is linked to smaller vesicular structures.

**FIGURE 4.**
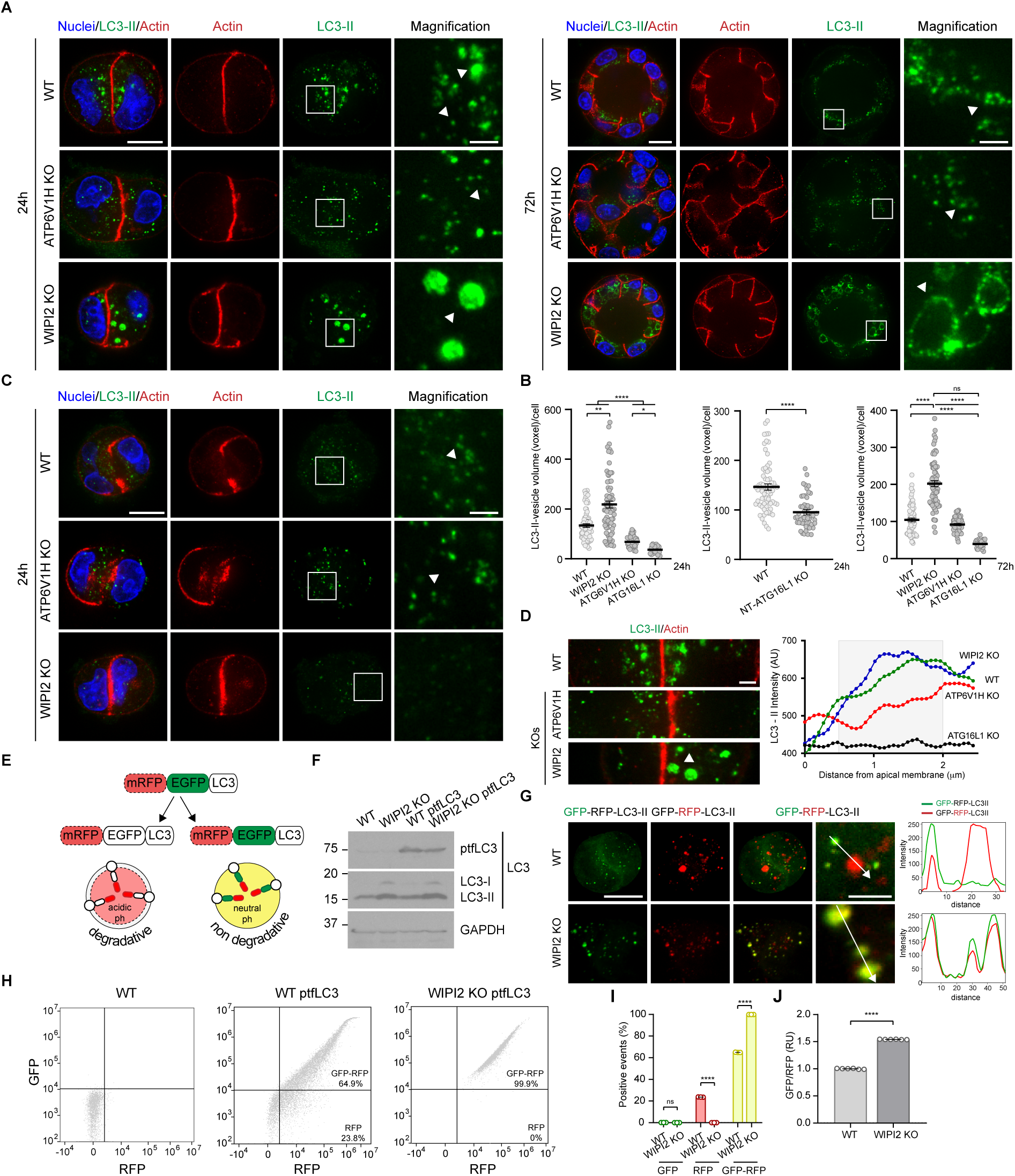
Noncanonical autophagy is mediated by large, non-degradative, LC3-positive vesicles in MDCK cysts. **(A)** 24-72h cysts show large LC3-positive vesicles in WIPI2 KO, large and small vesicles in WT, and small vesicles in ATP6V1H KO, after chloroquine (CQ) treatment. **(B)** Quantification of LC3-positive vesicle volume in WT, WIPI2 KO, ATP6V1H KO, and ATG16 KO cysts after CQ treatment at 24 (N=217 cyst) and 72h (N=224 cysts) of development. Kruskal-Wallis test was performed. Mann-Whitney test was conducted to analyze LC3-positive vesicle volume in NT-ATG16L1 KO 24h cysts (N=121 cysts). **(C)** IF of 24h cysts show the disappearance of large LC3-positive vesicles in WT and WIPI2 KO upon Bafilomycin A1 treatment. **(D)** LC3-positive vesicles in WT and WIPI2 KO show apical localization compared to ATP6V1H KO, upon CQ treatment in 24h cyst (left). LC3-II intensity quantified from the apical membrane in the images shown in (A) (right). **(E)** Schematic diagram of mRFP-EGFP-LC3 reporter to monitor autophagy. **(F)** WB of LC3 expression in WT and WIPI2 KO stable ptfLC3 cell lines. **(G)** Degradative (red) and non-degradative (yellow) vesicles in WT and WIPI2 KO 24h cysts (left). Fluorescence intensity profile of degradative (red) and non-degradative (red and green) are plotted along the white lines (right). **(H)** FACs dot blot representation of GFP and mRFP distribution in WT, WT ptfLC3, and WIPI2 KO ptfLC3. **(I)** Positive events for GFP, mRFP, and mRFP-GFP/ MFI values quantified by FACs in 24h cysts from WT ptfLC3 and WIPI ptfLC3. Data are expressed as the mean ± SEM from 3 independent experiments. Two-way ANOVA test was performed. **(J)** WIPI2 KO cysts show elevated GFP/RFP ratio compared to WT. Relative units (RU). Data are expressed as the mean ± SEM from 6 independent experiments. Unpaired two-tailed t-test was conducted. In panels A, C, and G, scale bars=10 μm and 2μm (magnification). In panels A and C, nuclei are stained with DAPI (blue). *P* values= \*\*\*\**p*<0.0001; \*\**p*<0.01, and \**p*<0.05; ns=non-significant.

When assessing vesicle localization, we observed that LC3-positive vesicles in WIPI2 KO and WT cysts showed a pronounced apical bias and asymmetric cytoplasmic distribution, especially evident at the 24-hour stage (Figure 4D). Conversely, ATP6V1H KO cysts showed a more uniform and dispersed vesicle distribution throughout the cytoplasm. The apical enrichment of LC3-positive vesicles specifically in the context of noncanonical autophagy may indicate a spatially restricted function in supporting or remodeling luminal architecture.

Canonical autophagy is primarily degradative, whereas noncanonical autophagy can include both degradative and non-degradative functions depending on the cellular context^16,17^. Based on this functional divergence, we next examined whether these vesicle populations differed in their degradative capacity. For this, we employed the tandem fluorescent-tagged LC3 (ptfLC3) biosensor, in which LC3 is fused to both GFP and mRFP (Figure 4E). This system takes advantage of the pH sensitivity of GFP, which is quenched in acidic lysosomes, whereas mRFP remains stable^49^. As such, degradative compartments (e.g., autolysosomes) appear red, while non-degradative vesicles—where both fluorophores remain intact— appear yellow. We generated stable ptfLC3-expressing cell lines in WT and WIPI2 KO backgrounds. In both, ptfLC3 expression recapitulated endogenous LC3 patterns (Figure 4F). After 24 hours of cyst development, we analyzed ptfLC3-labeled vesicles by immunofluorescence, measuring intensity variations across vesicles (Figure 4G). WT cysts displayed a combination of red and yellow vesicles, indicating the coexistence of degradative and non-degradative structures. In contrast, WIPI2 KO cysts exhibited predominantly yellow vesicles. This was further supported by flow cytometry, which revealed that nearly all vesicles in WIPI2 KO cells were yellow, in contrast to the mixed red/yellow distribution observed in WT cells (Figures 4H–4I). WIPI2 KO cells showed increased GFP fluorescence and a reduction in red-only (mRFP) vesicles compared to WT. The elevated GFP/RFP ratio in these cells indicates that LC3-positive vesicles are less acidified and thus not progressing to full lysosomal degradation (Figure 4J).

Taken together, these findings reveal that noncanonical autophagy is mediated by large, non-degradative, LC3-positive vesicles that are spatially biased toward apical regions and do not undergo lysosomal acidification. These features distinguish noncanonical autophagy structurally and functionally from canonical autophagy, and further support its specialized role in epithelial lumen resolution.

### Canonical and Noncanonical Autophagy Are Differentially Regulated to Support Lumenogenesis and Homeostasis

Our findings thus far demonstrate a functional divergence between canonical and noncanonical autophagy during lumen formation. Hence, we hypothesized that the balance between these two pathways fluctuates dynamically under basal conditions and may vary depending on the cellular process or metabolic state. These fluctuations may reflect a finely tuned mechanism that ensures proper lumen development and homeostasis. To this end, we analyzed the autophagic flux in WT cells undergoing lumenogenesis upon treatment with BAF and CQ. Autophagic activity was assessed at six distinct time points, ranging from 3 to 72 hours, thereby encompassing the entire process (Figure 5A).

**FIGURE 5.**
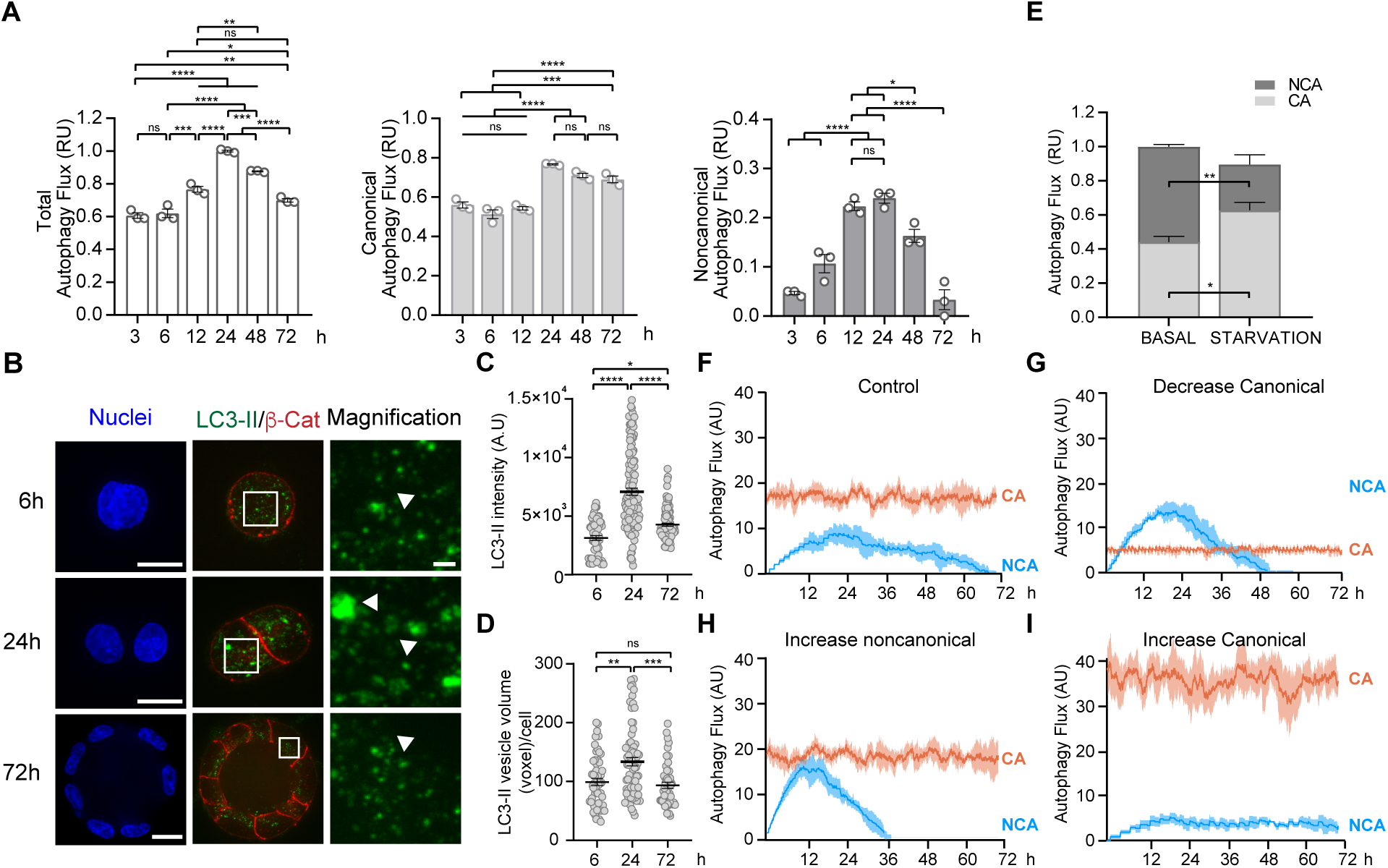
Canonical and noncanonical autophagy pathways are dynamically regulated during epithelial morphogenesis. **(A)** Total and noncanonical autophagic flux (AF) peak at 24h, whereas canonical autophagy remains relatively stable throughout cyst development. AF was assessed by quantifying LC3-II MFI via FACs throughout cyst development. Data are expressed as the mean ± SEM from 3 independent experiments. One-way ANOVA test was performed. **(B)** 24h cysts show large LC3-positive vesicles. Nuclei are stained with DAPI (blue). Scale bars=10 μm and 1 μm (magnification). **(C-D)** LC3-II intensity and the volume of LC3-II–positive vesicles are increased in 24h cysts, as determined by quantitative analysis of the IF images shown in panel (B). Data are expressed as the mean ± SEM from 3 independent experiments. (N=280 and 169 cysts, respectively). Kruskal-Wallis test was performed. **(E)** Under 3h of starvation canonical AF increases, whereas noncanonical AF decreases in 24h cysts. Data are expressed as the mean ± SEM from 3 independent experiments. Two-way ANOVA t-test was conducted. CA: canonical autophagy, NCA: noncanonical autophagy. **(F-I)** CA and NCA pathways are interconnected. Graphics based on an agent-based model predict how fluctuation in one pathway affect the other. The four conditions include: normal conditions (F), decreased CA (G), increased NCA (H), and increased CA (I). In panels A and E, Relative units (RU). In panels C, F-I, Arbitrary units (AU). *P* values= \*\*\*\**p*<0.0001; \*\*\**p*<0.001; \*\**p*<0.01; \**p*<0.05; ns=non-significant.

Under basal conditions, total autophagic flux progressively increased, peaking at 24 hours, followed by a gradual decline. This temporal profile was confirmed by immunofluorescence analysis of LC3-positive vesicle intensity, which reaffirmed a peak at 24 hours of cyst development (Figures 5B–5C). When canonical and noncanonical autophagy were assessed independently, we observed that canonical autophagy was the predominant pathway, exhibiting a flux approximately twice as high as noncanonical autophagy (Figure 5A). Nonetheless, while canonical flux remained relatively stable over time, noncanonical autophagy showed a sharper increase at 12 and 24 hours, followed by a more marked decline thereafter (Figure 5A). The peak in noncanonical flux correlated with an increased presence of large LC3-positive vesicles (Figure 5D), similar to those observed in WIPI2 KO cells, previously linked to noncanonical autophagy (Figure 4). These results suggest that during early cyst development—particularly as polarity is being established and extensive remodeling of apical and basolateral surfaces occurs—noncanonical autophagy is transiently elevated. In contrast, canonical autophagy provides a more constant baseline level of activity, potentially serving housekeeping and homeostatic roles.

To further elucidate how the balance between canonical and noncanonical autophagy adapts to environmental stress, we exposed WT cysts to nutrient deprivation—a condition known to stimulate canonical autophagy^50,51^. Under starvation, cyst development halted (Figure S5A) while a single, significantly larger lumen formed (Figures S5B–S5D). This observation can be explained by the ceasing of cellular division despite ongoing canonical autophagy, allowing the lumen to expand without an increase in spheroid volume. Notably, canonical autophagy flux increased by nearly 50%, whereas noncanonical flux decreased by a similar margin (Figure 5E and S5E), suggesting that under metabolic stress, canonical autophagy is prioritized at the expense of noncanonical activity

Because canonical and noncanonical autophagy share fundamental molecular machinery, we hypothesized that some level of interdependence must exist. To test this, we developed an agent-based model simulating autophagy dynamics in epithelial morphogenesis. This model incorporated experimental data from WT cysts and allowed us to predict how fluctuations in one pathway may affect the other. Simulations recapitulated the canonical/noncanonical autophagy balance observed under basal conditions (Figure 5F), as well as experimentally induced scenarios in which either pathway was suppressed or elevated (Figures 5G–5I). Model predictions showed that under normal conditions, noncanonical autophagy increases rapidly, peaks at around 24 hours, and returns to baseline by 72 hours—without significantly altering canonical activity (Figure 5F). However, when canonical autophagy was reduced, as observed in WIPI2 KO cysts (Figure 3C), noncanonical autophagy reached higher maximum levels (Figure 5G). In contrast, an artificial increase in canonical activity suppressed noncanonical flux (Figure 5I), whereas increasing noncanonical activity did not affect canonical autophagy rates (Figure 5H). These predictions were consistent with our experimental data from ATP6V1H KO cysts and nutrient-deprived WT cysts (Figures 5E and S5E).

In conclusion, our findings reveal that canonical and noncanonical autophagy pathways are dynamically regulated during epithelial morphogenesis and fulfill distinct yet complementary roles. Canonical autophagy provides a stable baseline of degradative activity critical for cellular homeostasis and nutrient recycling, particularly under metabolic stress. In contrast, noncanonical autophagy is transiently activated during early stages of cyst development, supporting processes such as trafficking establishment and lumen resolution through large, non-degradative LC3-positive vesicles. These pathways are interconnected, showing reciprocal modulation depending on developmental timing and metabolic cues. While noncanonical autophagy can increase independently, canonical autophagy appears to suppress noncanonical activity when upregulated, suggesting a tightly coordinated balance between the two. Together, our experimental and computational analyses establish that the fine-tuned interplay between canonical and noncanonical autophagy is essential for orchestrating epithelial lumenogenesis and maintaining tissue architecture.

### Noncanonical Autophagy Orchestrates Membrane Remodeling During Epithelial Lumen Formation

Selective impairment of noncanonical autophagy disrupts single-lumen formation in cysts derived from single MDCK cell, resulting in a persistent multilumen phenotype. Nevertheless, in physiological contexts such as tubular organ development, epithelial structures emerge from multicellular clusters that first form small mini-lumens, which later coalesce into a single, continuous lumen through a process known as lumen resolution^1,3^.

To model this more accurately, we generated MDCK epithelial tube micropatterns, a system that recapitulates epithelial morphogenesis in a spatially controlled and physiologically relevant context^52^. Compared to traditional MDCK cysts, epithelial tube micropatterns offer enhanced experimental power through precision microenvironmental control and reproducible architecture. We first assessed lumen formation in micropatterns derived from our autophagy-deficient cell lines (Figure 6A). As expected, localization of Podxl confirmed the successful establishment of a polarized epithelial axis and further quantitative analysis of lumen formation revealed patterns consistent with our cyst-based models (Figures 2I–2J and 3G–3H): only WT and WIPI2 KO cells, both of which retain noncanonical autophagy, successfully formed single lumens (Figures 6A–6B). Consistent with our previous observations in cysts (Figures 3G– 3I), epithelial tubes deficient in WIPI2 displayed impaired lumen expansion (Figure 6A), which may be linked to reduced Claudin-2 expression (Figure 6C)—a phenomenon also observed in cysts (Figures S4H–S4I).

**FIGURE 6.**
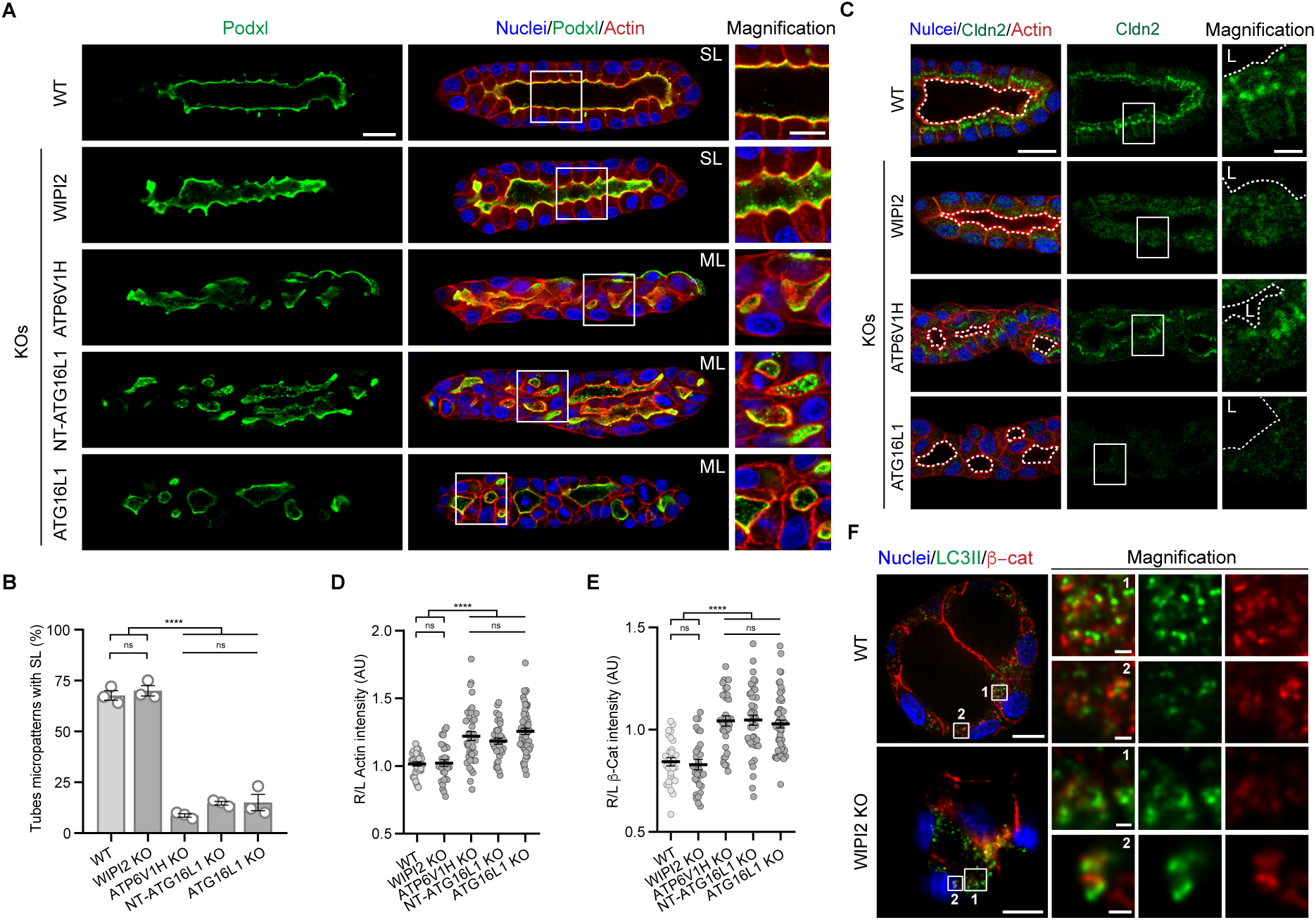
Noncanonical autophagy-deficient cells fail to remodel membranes effectively, resulting in multilumen epithelial tubes. **(A)** Epithelial tubes lacking noncanonical autophagy (ATP6V1H1 KO, NT-ATG16L1 KO, and ATG16L1 KO) fail to resolve multi lumens (ML) into a single lumen (SL). Epithelial tubes lacking canonical autophagy (WIPI2 KO) show impaired lumen expansion. IF of 96h epithelial tubes derived from WT, WIPI2 KO, ATP6V1H KO, NT-ATG16L1 KO, and ATG16L1 KO cells. Podocalyxin (Podxl). Scale bars= 20 μm and 10 μm (magnification). **(B)** Percentage (%) of SL quantified from images shown in (A). Data are expressed as the mean ± SEM from 3 independent experiments (N=1500 epithelial tubes). One-way ANOVA test was performed. **(C)** Epithelial tubes lacking canonical autophagy (WIPI2 KO and ATG16L1 KO) exhibit reduced levels of Claudin-2 (Cldn2). Scale bars=20 μm and 5 μm (magnification). **(D-E)** Actin and β-catenin (β-cat) accumulate at remodeling membranes in epithelial tubes lacking noncanonical autophagy (ATP6V1H1 KO, NT-ATG16L1 KO, and ATG16L1 KO). Quantitative analysis shows the fluorescence intensity ratio of actin and β-cat between remodeling (R) and lateral (L) membranes. Data are represented as the mean ± SEM from 3 independent experiments. One-way ANOVA was performed. **(F)** β-cat is trafficked to large, LC3-positive vesicles. Images shows co-localization of LC3-II and β-cat at vesicle structures in WT and WIPI2 KO. Scale bars =10 μm and 1 μm (magnification). In panels A, C and F, nuclei are stained with DAPI (blue.). In panels D and E, Arbitrary units (AU). *P* values= \*\*\*\**p*<0.0001; ns=non-significant.

To further validate the requirement for noncanonical autophagy in lumen resolution, we took advantage of a density-dependent feature of MDCK cyst morphogenesis. At very low plating densities, MDCK cells predominantly form clonal cysts through cell proliferation. Nevertheless, at high cell densities, both proliferation and aggregation occur simultaneously, leading to the formation of isolated mini-lumens that subsequently fuse into a single lumen^53^. Quantitative analysis confirmed an inverse correlation between plating density and initial single-lumen formation in WT cysts (Figure S6A). Under low-density conditions, the majority of cysts formed single lumens by 72 hours. In contrast, high-density cultures showed increased multilumen structures at the same time point. Notably, by 96 hours, most high-density WT cysts resolved into single lumens, matching the efficiency observed in low-density conditions at 72 hours. High-density WT cysts also displayed enlarged LC3-positive vesicles—reminiscent of those observed in WIPI2 KO cells (Figure S6B)—a morphological hallmark of noncanonical autophagy. The observed upregulation of noncanonical autophagy may reflect a cellular mechanism aimed at resolving the ectopic lumens induced by the high cell density. In agreement with our earlier results, only WIPI2 KO cysts, which maintain noncanonical autophagy, retained the ability to resolve into single lumens. Cysts derived from ATG16L1 KO, NT-ATG16L1 KO, or ATP6V1H KO cells lacked this capacity (Figure S6A).

Aiming to investigate the mechanistic involvement of noncanonical autophagy in lumen resolution, we then focused on two converging observations: (1) cells deficient in the noncanonical autophagy pathway fail to resolve multilumen structures into a single central lumen; and (2) under basal conditions, noncanonical autophagy exhibits a sharp, transient peak at 24 hours—precisely during the window of active membrane remodeling associated with lumen coalescence. These findings suggest that noncanonical autophagy, through the formation of large LC3-positive, non-degradative vesicles, plays a direct role in orchestrating membrane remodeling events required for converting mini-lumens into a single, continuous epithelial lumen during morphogenesis.

Actin and β-catenin are essential regulators of membrane organization in epithelial cells. Branched actin networks promote directional cytoskeletal remodeling, while β-catenin connects E-cadherin to the actin cytoskeleton, supporting junctional stability. Their accumulation at remodeling sites can signal disruption of trafficking or turnover processes, potentially interfering with proper lumen formation^3,54^. To determine whether deficiencies in noncanonical autophagy alter the dynamics and distribution of actin and β-catenin during epithelial tube formation and in high-density MDCK cysts, we conducted a series of experiments. Our analysis of epithelial tubes revealed that cells lacking CASM exhibited an approximate 20% increase in actin and β-catenin signals at sites of membrane remodeling, when compared to WT controls. Furthermore, high-density MDCK cysts derived from noncanonical autophagy-deficient lines (ATG16L1 KO and ATP6V1H KO) showed a marked accumulation of both proteins in remodeling membranes, with actin increasing by roughly 10% and β-catenin by over 30% (Figures 6D-6E and S6C–S6F).

We then evaluated β-catenin trafficking by performing biotinylation assays, which confirmed its internalization in both WT and ATG16L1 KO cysts. However, only WT cells efficiently redistributed β-catenin, while CASM-deficient cysts retained it intracellularly for extended periods (Figures S6G–S6H). Complementary confocal analysis showed that in WT cysts, β-catenin was internalized and trafficked to large, LC3-positive vesicles. This effect was even more pronounced in WIPI2 KO cysts, further supporting a role for noncanonical autophagy in junction remodeling (Figure 6F). To gain deeper insight into this mechanism, we inhibited endocytic trafficking using Dynasore—a dynamin inhibitor that also blocks recycling pathways. Under these conditions, high-density WT cysts failed to form a single lumen (Figure S6I), indicating that endocytic remodeling is critical for lumen resolution. Overall, these findings support a model in which the V-ATPase/ATG16L1-dependent noncanonical autophagy pathway (CASM) orchestrates the endocytic and recycling events required for membrane reorganization during lumen formation.

### The CASM Pathway Coordinates Endocytic Recycling and Membrane Remodeling During Lumen Formation

Membrane remodeling is a highly regulated process involving endocytosis, recycling, sorting, and degradation. Through endocytosis, membrane proteins and lipids are internalized and either targeted for degradation or returned to the membrane via recycling^55,56^. This system is largely orchestrated by Rab GTPases and actin/microtubule dynamics. Rab5 regulates early endosome formation, Rab7 governs late endosome formation, and Rab11 is essential for membrane recycling^12,57,58^.

Within this context, a substantial reduction in Rab11-positive vesicles was detected in CASM-deficient cysts (ATG16L1 KO, ATP6V1H KO, and NT-ATG16L1 KO), suggesting impaired recycling (Figures 7A–B). In contrast, WIPI2 KO cysts, which lack canonical autophagy but retain noncanonical activity, showed a non-significant trend toward apical Rab11 accumulation. Additionally, the localization and abundance of Rab5 were diminished in CASM-deficient cysts, while remaining largely unchanged in canonical-autophagy-deficient models (Figures 7C-7D).

**FIGURE 7.**
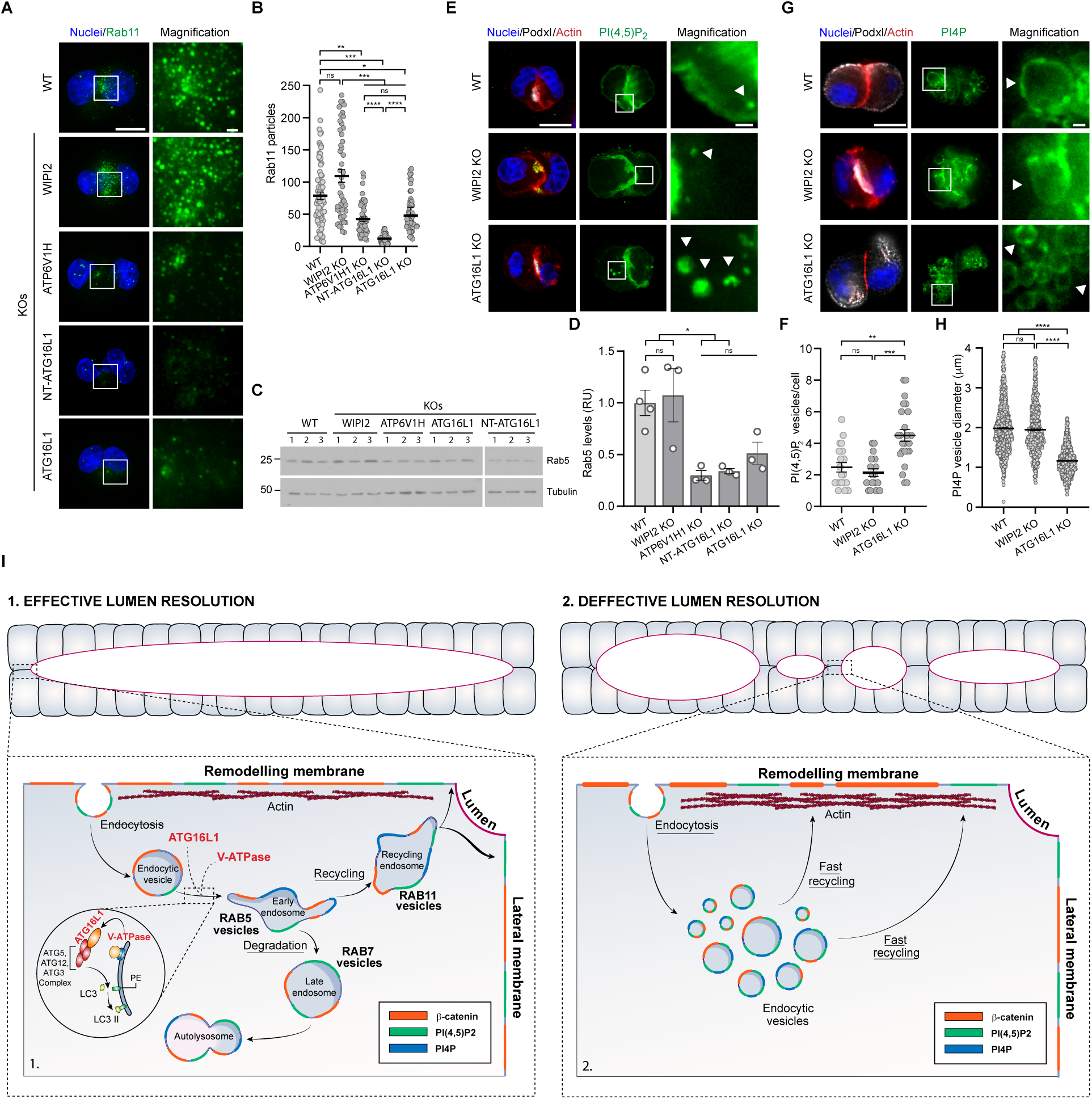
Noncanonical autophagy supports endocytic recycling during epithelial lumen formation. **(A)** Rab11 positive vesicles are reduced in noncanonical deficient 24h cysts (ATP6V1H KO, NT-ATG16L1 KO, and ATG16L1 KO). Scale bars=20 μm and 1 μm (magnification). **(B)** Rab11–positive vesicles quantified in IF images shown in (A). Data are expressed as the mean ± SEM from 3 independent experiments. (N=297 cells). Kruskal-Wallis test was performed. **(C)** Noncanonical defective cysts (ATP6V1H KO, NT-ATG16L1 KO, and ATG16L1 KO) exhibit reduced Rab5 levels, assessed by WB. **(D)** Rab5 levels quantified in WB shown in (C). One-way ANOVA test was performed. **(E)** PI(4,5)P_2_ accumulate in intracellular vesicles in 24h cysts derived from ATG16L1 KO. Scale bars=10 μm and 5 μm (magnification). Relative units (RU). **(F)** PI(4,5)P_2_ vesicles quantification from IF images in (E). Data are expressed as the mean ± SEM from 3 independent experiments. (N=61 cells). Kruskal-Wallis test was performed **(G)** PI4P vesicles were increased in number and reduced in size in 24h cysts derived from ATG16L1 KO. Scale bars=10 μm and 1 μm (magnification). **(H)** PI4P vesicles quantification from IF images in (G). Data are expressed as the mean ± SEM from 3 independent experiments. (N=3, 3102 vesicles from 114 cells). Kruskal-Wallis test was performed. *P* values= \*\*\*\**p*<0.0001; \*\*\**p*<0.001; \*\**p*<0.01, and \**p*<0.05; ns=non-significant. (**I**) Graphical model showing the effect of canonical and noncanonical CASM autophagy in lumen resolution of epithelial tubes.

Given that endosomal identity and function are tightly controlled by phosphoinositide composition, we next examined the distribution of key phosphoinositide species [PI(4,5)P₂, PI4P, PI(3,4)P₂, and PI(3,4,5)P₃]. PI(4,5)P₂ is enriched at the plasma membrane and early endosomes, where it facilitates clathrin-mediated endocytosis and membrane curvature^6^. Our analysis revealed a marked accumulation of PI(4,5)P₂ in intracellular vesicles of CASM-deficient cysts (Figure 7E-7F), suggesting a disruption in endosomal lipid dynamics. Notably, this phenotype was absent in WIPI2 KO cells, indicating that the defect is specific to noncanonical autophagy rather than a general autophagy impairment. By contrast, PI(3,4,5)P₃ and PI (3,4)P_2_ were mostly unaffected in CASM-deficient and canonical defective cyst, suggesting that basolateral membrane identity was not compromised in these cells (Figure S7A).

During early endosomal maturation, PI(4,5)P₂ is converted to PI4P, enabling V-ATPase recruitment. In particular, the V-ATPase subunit ATP6V0A2 binds directly to PI4P through its cytosolic domain, a brief but critical interaction that anchors the V-ATPase to endosomal membranes and drives the progression of endosomal maturation^59^. Indeed, we observed that PI4P-decorated endocytic vesicles were significantly reduced in size, while increased in number, in CASM-deficient cells compared to controls, suggesting that endosome maturation is severely compromised (Figure 7G-7H). These findings suggest that CASM deficiency hinders early endosomal maturation by disrupting the phosphoinositide transitions necessary for V-ATPase recruitment and proper recycling, ultimately affecting endosomal function and membrane remodeling.

Finally, we characterized the effect of autophagy defect in spindle orientation. Proper spindle orientation is crucial for maintaining epithelial tissue integrity and ensuring that daughter cells inherit correct polarity and remain within the monolayer^1,3^. Aberrant endocytosis disrupts the spatial cues or membrane composition required for correct spindle positioning, leading to misoriented divisions and tissue disorganization^60^. Notably, we observed spindle orientation defects exclusively associated with noncanonical autophagy deficiency. These defects are likely a direct consequence of an impaired membrane remodeling mechanism that arises under conditions of noncanonical autophagy disruption (Figures S7B-S7D).

Together, these results establish that noncanonical autophagy, through the CASM pathway, plays a central role in orchestrating membrane remodeling during epithelial lumen formation (Figure 7I and S7E). CASM supports the endocytic recycling machinery required to internalize and redistribute junctional and cytoskeletal components, including β-catenin and actin, and regulates the maturation of endosomes through Rab GTPases and phosphoinositide remodeling. Disruption of CASM leads to defective trafficking, impaired junctional remodeling, and failure to resolve multiple lumens into a single central cavity. These findings position CASM as a fundamental mechanism driving lumen morphogenesis through precise control of membrane dynamics.

## DISCUSSION

In this study, we uncover a pivotal role for noncanonical autophagy—specifically the CASM (Conjugation of ATG8 to Single Membranes) pathway—in orchestrating membrane remodeling and lumen morphogenesis during epithelial development. Using a combination of huNTOrgs, MDCK cysts, and epithelial tube micropatterns, we demonstrate that ATG16L1, a core autophagy protein, is essential for proper lumen resolution. While canonical autophagy alone is insufficient to rescue the multilumen phenotype observed in ATG16L1-deficient organoids and cysts, restoring noncanonical autophagy reinstates the ability to form a single, central lumen (Figure 7I). Through systematic dissection of ATG16L1 functional domains and generation of pathway-specific KO models (targeting ATP6V1H for noncanonical autophagy and WIPI2 for canonical), we show that canonical autophagy supports lumen expansion and junctional integrity, whereas noncanonical autophagy is required for resolving mini-lumens into a unified central cavity. Mechanistically, we identify that CASM operates through a large LC3-positive vesicle population that is non-degradative and localized apically—distinct from canonical autophagic compartments. These vesicles are enriched during key morphogenetic stages and fail to mature properly when CASM is impaired. Loss of CASM disrupts β-catenin and actin dynamics, impairs endocytic recycling, and alters the localization of Rab GTPases (Rab5, and Rab11), as well as phosphoinositide composition critical for endosomal maturation. We further show that CASM-deficient cells accumulate PI(4,5)P₂ and altered PI4P vesicle size and distribution, suggesting defects in the lipid transitions that prime endosomes for acidification and recycling. Computational modeling supports a dynamic, reciprocal regulation between canonical and noncanonical autophagy, with CASM becoming transiently elevated during early cyst development and suppressed under nutrient-limiting conditions where canonical autophagy is prioritized. Altogether, our findings define noncanonical autophagy not as a passive or compensatory pathway, but as an active, instructive mechanism that coordinates membrane remodeling events necessary for epithelial lumen formation and morphogenesis (Figure 7I and S7E).

These findings offer significant insights into the functional complexity of autophagy during epithelial morphogenesis, particularly in establishing and resolving epithelial lumens. While canonical autophagy has been extensively studied in the context of nutrient stress and homeostasis^50,51^, its role in tissue architecture has remained ambiguous. Our results clarify that canonical autophagy is primarily supportive, promoting junctional stability and contributing to lumen expansion, but it is not sufficient to resolve multiple lumens into a single, continuous cavity. Instead, noncanonical autophagy—via the CASM pathway—emerges as an active, mechanistically distinct contributor to morphogenesis, coordinating membrane remodeling and endocytic recycling in a spatially and temporally restricted manner. This distinction is critical, as it elevates noncanonical autophagy from a peripheral or redundant process to a central regulator of morphogenetic events.

Our findings also intersect with a growing body of literature implicating noncanonical autophagy in membrane remodeling beyond degradation. For instance, LC3-associated phagocytosis (LAP), LC3-associated endocytosis (LANDO), and CASM share a common molecular framework involving ATG8 lipidation at single membranes independent of the canonical initiation complex^61,62^. Recent work has shown that CASM is recruited at sites of membrane repair and endocytic maturation, particularly in contexts involving V-ATPase activation and Rab-dependent trafficking^63–65^. Our demonstration that CASM-deficient cells accumulate PI(4,5)P₂, and exhibit disrupted Rab11 localization suggests that noncanonical autophagy is not simply decorating membranes with LC3, but is actively coordinating vesicle identity and fate. These processes are fundamental to lumen resolution, where precise trafficking, polarity, and membrane removal are essential.

The ability of noncanonical autophagy to control these events has potential implications beyond organoid models. During embryonic development, secondary neurulation and epithelial tube formation rely on tightly regulated morphogenetic events that require the rapid formation and reorganization of apical domains^3,7,53^. The failure of ATG16L1-deficient huNTOrgs to resolve into a single central lumen may provide insight into developmental disorders such as spina bifida or other neural tube defects, where epithelial closure and lumen resolution are impaired. Moreover, given the involvement of noncanonical autophagy in immune responses and epithelial repair^21,66^, these findings may extend to regenerative contexts and disease models involving barrier dysfunction or aberrant epithelial remodeling.

Looking ahead, the identification of large, non-degradative LC3-positive vesicles as central components of the CASM response opens exciting avenues to explore their cargo, interactors, and biogenesis. Future studies using proximity labeling, live-cell imaging, and proteomic analysis could reveal how these vesicles selectively coordinate recycling versus degradation and how they interface with endosomal maturation machinery. Understanding how CASM is activated and terminated, and how it integrates with signaling pathways such as WNT, Notch, or BMP, could also provide further insight into its developmental and pathological roles.

While our findings provide strong evidence for a central role of noncanonical autophagy in epithelial lumen formation, several limitations should be acknowledged. The in vitro systems used, including MDCK cysts, epithelial micropatterns, and organoids, do not fully capture the complexity of in vivo tissue architecture or signaling environments. Additionally, although our genetic models effectively dissociate canonical from noncanonical autophagy, the shared core machinery between the pathways may introduce overlapping effects or compensatory responses. Our imaging-based analysis of endocytic remodeling was limited to fixed-cell observations and endpoint measurements, precluding real-time tracking of vesicle dynamics. Finally, while our computational model offers valuable insight into autophagy regulation, it simplifies complex cellular interactions and requires further validation in vivo. Future work using live imaging, proteomics, and in vivo models will be critical to expand and refine our outcomes.

In conclusion, our work establishes noncanonical autophagy as a fundamental driver of epithelial lumenogenesis, redefining its role from degradative support to a spatially regulated, mechanistically unique membrane remodeling system. These findings not only illuminate a previously underappreciated layer of morphogenetic regulation but also provide a framework for investigating CASM in development, disease, and regenerative biology.

## ACKNOWLEDGMENTS

We thank C. M. Ruiz-Jarabo for her comments on the manuscript. We thank members of the Martin-Belmonte laboratory for helpful discussions. This work was supported by MICINN (PID2020-120367GB-I00; PID2023-151844OB-I00; RED2022-134927-T), Fondos FEDER investigación (Horizon-MSCA-2022-DN-101119504) and Fundación Ramon Areces (CIVP18A3904). HR22-00447 from äLa Caixä, RTI2018-101586-A-I00, PID2021-126298OB-100 to N.M.-M. G.B. was supported by a FPI grant from MINECO (CSIC). T.A. was supported by by a Juan de la Cierva grant from MICINN, H2020-MSCA-IF-2019-897948 and Ayuda Investigador from AECC. G.H. was supported by a FPI grant from MINECO (BES-2018-077789. We are grateful to the Advance Microscopy, Flow cytometry and Animal facilities at the CBMSO. We also want to thank Dr. D. Bryant (University of Glasgow, The United Kingdom) and Dr F. Pimentel (CBM, Madrid, Spain) for the generous donation of PIP-probe plasmids, and ATG16L1 domain constructs.

---

**FIGURE S1.**
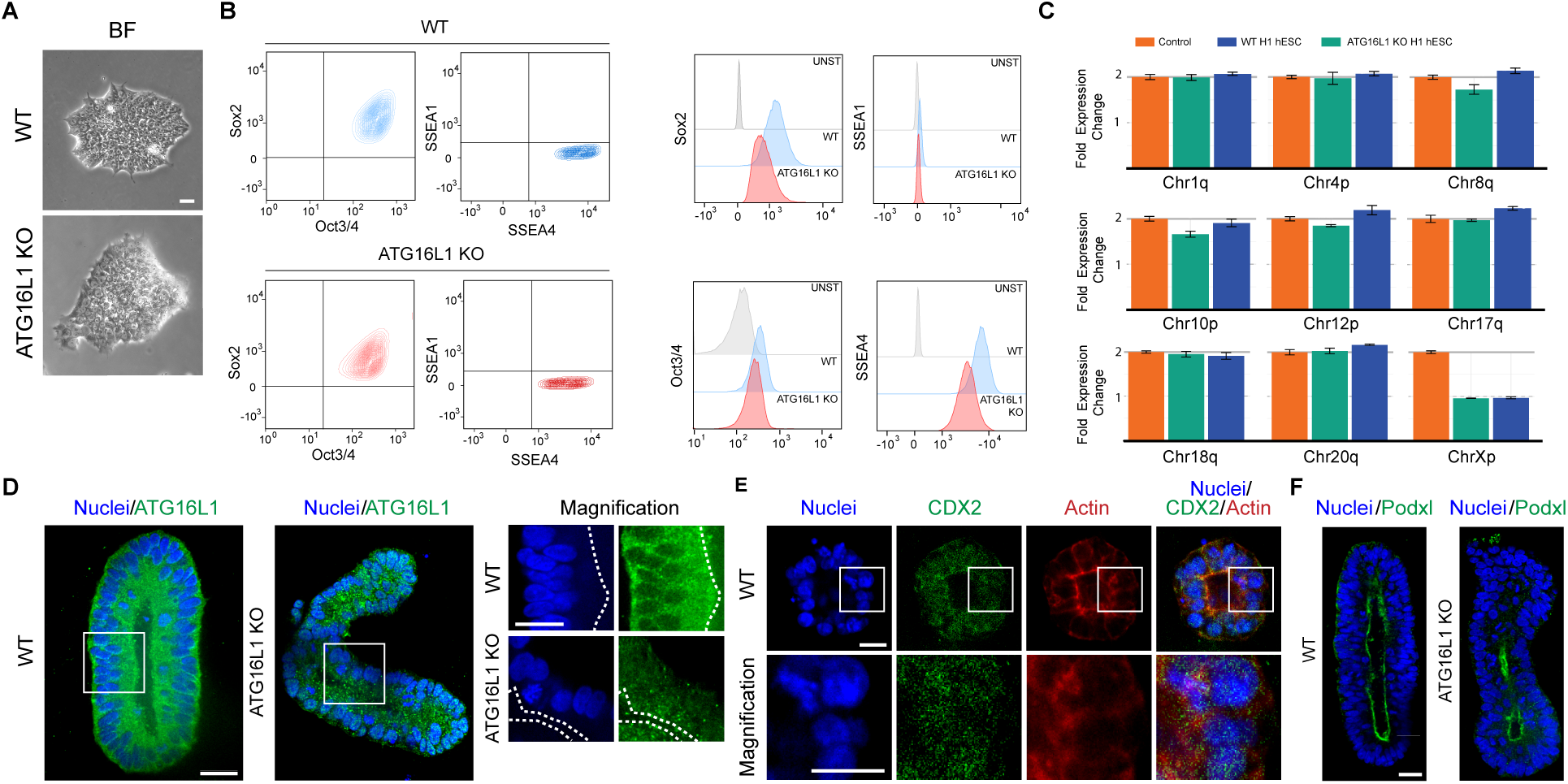
ATG16L1 KO H1 human Embryonic Stem Cells (hESCs) maintain a pluripotent state and preserve normal karyotype. **(A)** Bright-field images of WT and ATG16L1 KO H1 hESCs colonies showing undifferentiated morphology. Scale bar=10 μm. **(B)** WT and ATG16L1 KO H1 hESCs maintain a pluripotent state. FACs dot blots representation of Sox2, Oct3/4, SSEA1, and SSEA4 distribution in WT and ATG16L1 KO H1 hESCs (left). Representative histogram showing the expression of Sox2, Oct3/4, SSEA1, and SSEA4 in WT and ATG16L1 KO hESCs (right). **(C)** WT and ATG16L1 KO H1 hESCs preserve a normal karyotype. **(D)** IF shows absence of ATG16L1 in ATG16L1 KO posterior human neural tube organoids (huNTOrgs) at day 5 of development. Scale bar=20 μm. **(E)** IF analysis showing expression of CDX2 in WT and ATG16L1 KO posterior huNTOrgs at day 2 of development. Scale bar=10 μm**. (F)** ATG16L1 KO huNTOrgs retain apical–basal polarity, as evidenced by apical Podocalyxin (Podxl) localization. Scale bar=20 μm. In panels D-F, nuclei are shown in blue.

**FIGURE S2.**
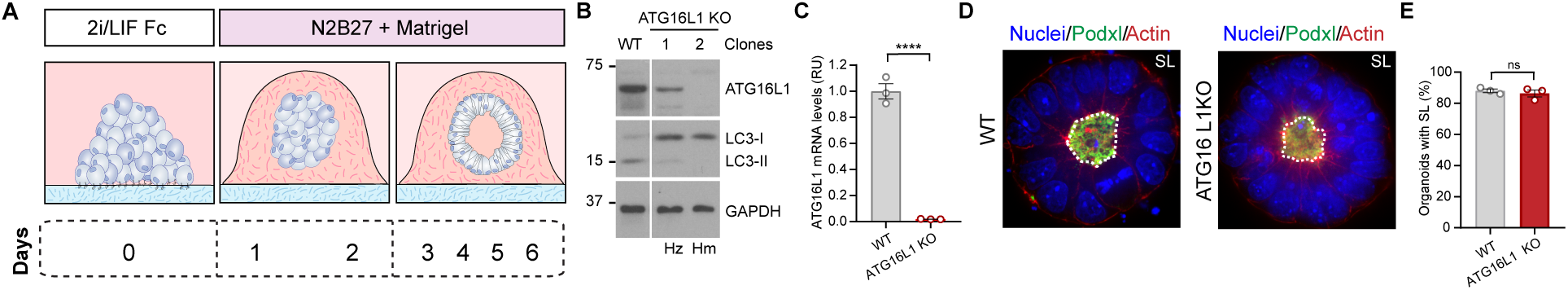
Autophagy is dispensable for lumen formation in organoids derived from mouse Embryonic Stem Cells (mESCs). **(A)** Schematic representation of the culture conditions used to model lumen formation in organoids derived from mESCs. **(B)** Absence of ATG16L1 expression impairs LC3 lipidation in ATG16L1 KO mESCs, assessed by WB. Hz: heterozygous. Hm: homozygous. **(C)** Quantification of ATG16L1 mRNA levels in ATG16L1 KO compare to WT mESCs. Data are expressed as the mean ± SEM from 3 independent experiments. Unpaired two-tailed t-test was conducted. **(D)** Autophagy deficient organoids derived from ATG16 KO mESCs form single lumens (SL). Nuclei are shown in blue. Scale bar=20 μm. **(E)** Percentage (%) of SL in organoids quantified from IF images in (D). Data are expressed as the mean ± SEM from 3 independent experiments. Unpaired two-tailed t-test was conducted. *P* value= \*\*\*\**p*<0.0001; ns=non-significant.

**FIGURE S3.**
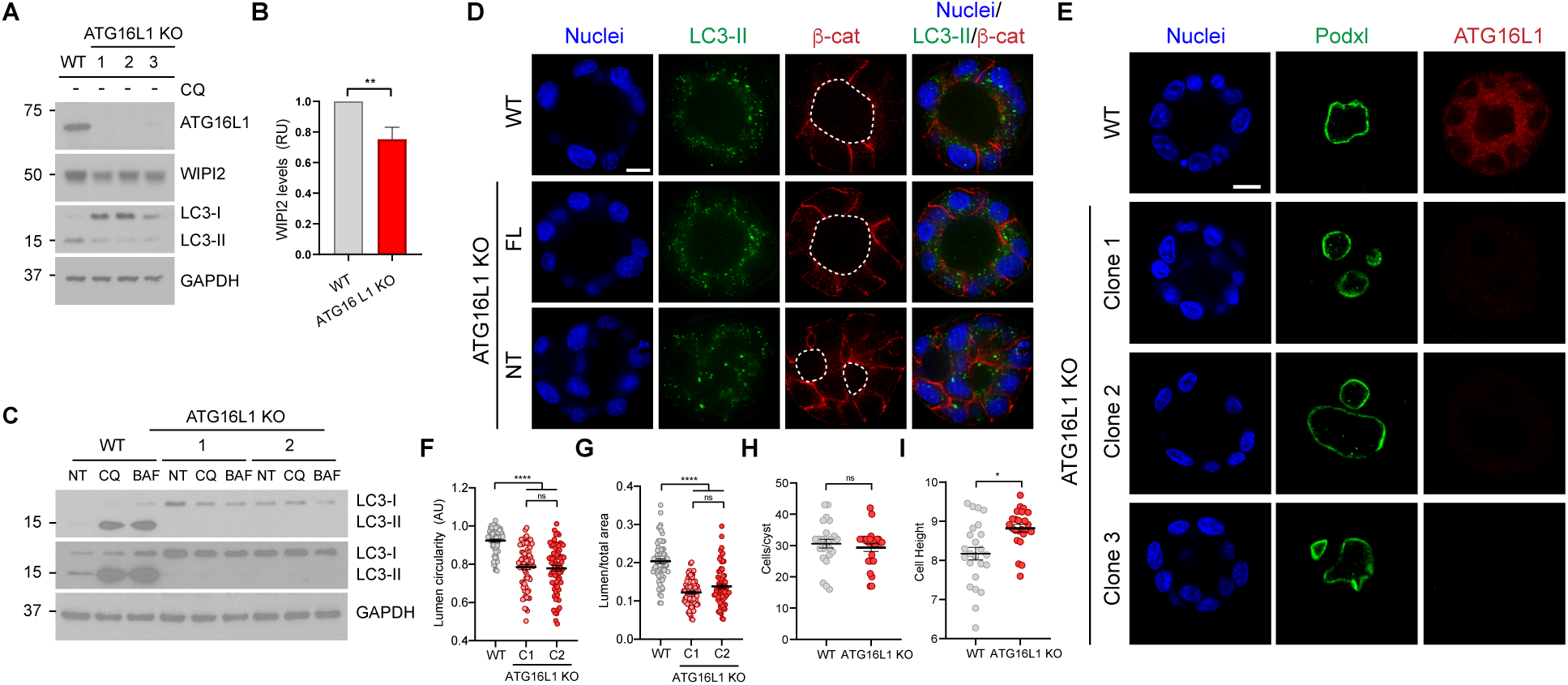
Impaired autophagy disrupts lumen resolution, reduces lumen area, and compromises lumen circularity in MDCK cysts. **(A)** Absence of ATG16L1 impairs LC3 lipidation and reduces WIPI2 protein levels in cysts derived from three ATG16L1 KO clones (1,2, and 3), assessed by WB. **(B)** Quantification of WIPI2 protein levels in WT and ATG16L1 KO WT cysts. Data are expressed as the mean ± SEM from 3 independent experiments. Unpaired two-tailed t-test was conducted. **(C)** Absence of ATG16L1 impairs LC3 lipidation in the absence (NT) or presence of Chloroquine (CQ) and Bafilomycin A1 (BAF) in cysts derived from two ATG16L1 KO clones (1 and 2), assessed by WB. **(D)** LC3 lipidation and lumen resolution in 72h cysts derived from WT and ATG16 KO cell lines stably expressing the full-length (FL) ATG16L1 or the NT-domain of ATG16L1. Scale bar=10 μm. **(E)** 72h cysts derived from three ATG16L1 KO cell lines (clones 1, 2, and 3) lack ATG16L1 expression and fail to resolve multiple lumens (ML) into a single lumen (SL), but still retain apical–basal polarity, as indicated by the apical localization of Podocalyxin (Podxl). **(F-G)** Quantitative analysis shows that 72h cysts with SL derived from two ATG16L1 KO cell lines (clones 1 and 2) exhibit reduced circularity and lumen area. Data are expressed as the mean ± SEM from 3 independent experiments. (N=204 cyst for lumen area, N=204 cysts for circularity). Kruskal-Wallis test was performed. **(H-I)** Quantitative analysis shows that 72h cysts with SL derived from ATG16L1 KO cells contain a similar number of cells but display increased cell height using CartoCell. Data are expressed as the mean ± SEM from 3 independent experiments. (N=50 cyst for cell number, N=50 cysts for cell height). Mann-Whitney test was performed. In panels D-E, nuclei are shown in blue. *P* values= \*\*\*\**p*<0.0001; \*\**p*<0.01, and \**p*<0.05; ns=non-significant.

**FIGURE S4.**
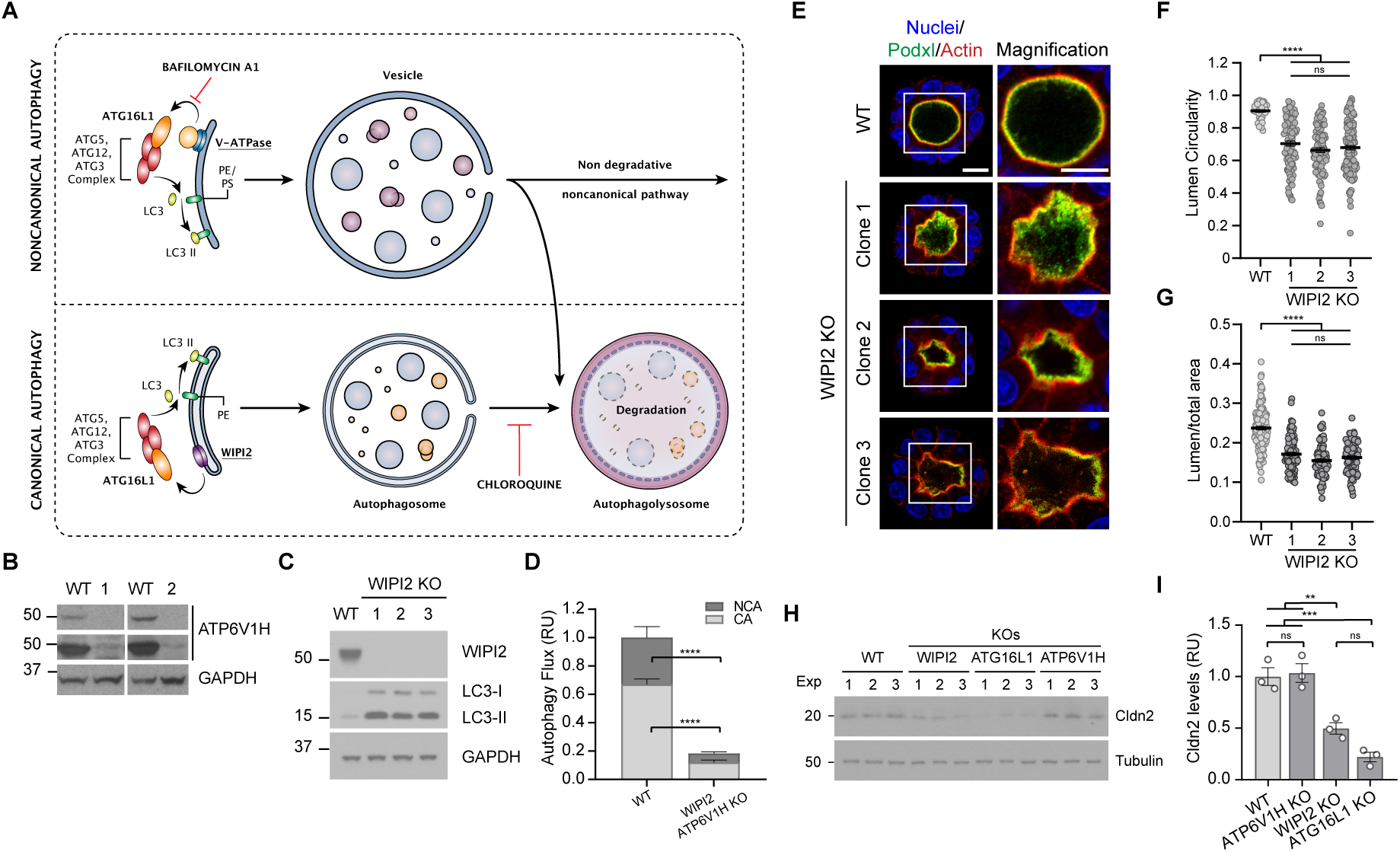
Canonical autophagy is essential for lumen expansion during epithelial morphogenesis in MDCK cysts. **(A)** Schematic model illustrating the noncanonical (degradative and non-degradative) and canonical (degradative) autophagy pathways. The vacuolar ATPase (V-ATPase) acts as a key regulator of noncanonical autophagy by directly interacting with ATG16L1 to facilitate LC3 recruitment to single-membrane compartments. WIPI2 plays a crucial role in canonical autophagy by recruiting ATG16L1 for targeting LC3 to double membrane compartments. Bafilomycin inhibits V-ATPase activity. Chloroquine impairs autophagosome fusion with lysosome. **(B)** Absence of ATP6V1H in ATP6V1H KO (clones 1 and 2) assessed by WB. **(C)** Absence of WIPI2 and LC3 lipidation in WIPI2 KO (clones 1, 2, and 3), assessed by WB. **(D)** WIPI2/ATP6V1H KO cysts exhibit an almost complete absence of autophagy. Autophagic Flux assessed by FACs, quantified by the mean fluorescence intensity of LC3-II, in 24h MDCK cysts from WT and WIPI2/ATP6V1H KO cells. CA: Canonical Autophagy Flux. NCA: Noncanonical Autophagy Flux. Data are expressed as the mean ± SEM from 5 independent experiments. Two-way ANOVA test was performed. **(E)** 72h cysts derived from WIPI2 KO clones present reduced lumen size. WIPI2 KOs cysts retain apical– basal polarity, as evidenced by apical Podocalyxin (Podxl) localization. Nuclei are shown in blue. Scale bar=10 μm. **(F-G)** Quantitative analysis shows that 72h cysts derived from WIPI2 KO clones exhibit reduced lumen circularity and smaller lumen area. Data are expressed as the mean ± SEM from 3 independent experiments. Kruskal-Wallis and one-way ANOVA tests were performed to analyze lumen circularity and lumen area data, respectively. **(H)** Claudin-2 (Cldn2) levels are reduced in 72h cysts derived from WIPI2 KO and ATG16L1 KO cells, both defective in canonical autophagy, assessed by WB. **(I)** Cldn2 levels quantification from the WB shown in panel (H). Data are expressed as the mean ± SEM from 3 independent experiments. One-way ANOVA test was performed. *P* values= \*\*\*\**p*<0.0001; \*\*\**p*<0.001; \*\**p*<0.01; ns=non-significant.

**FIGURE S5.**
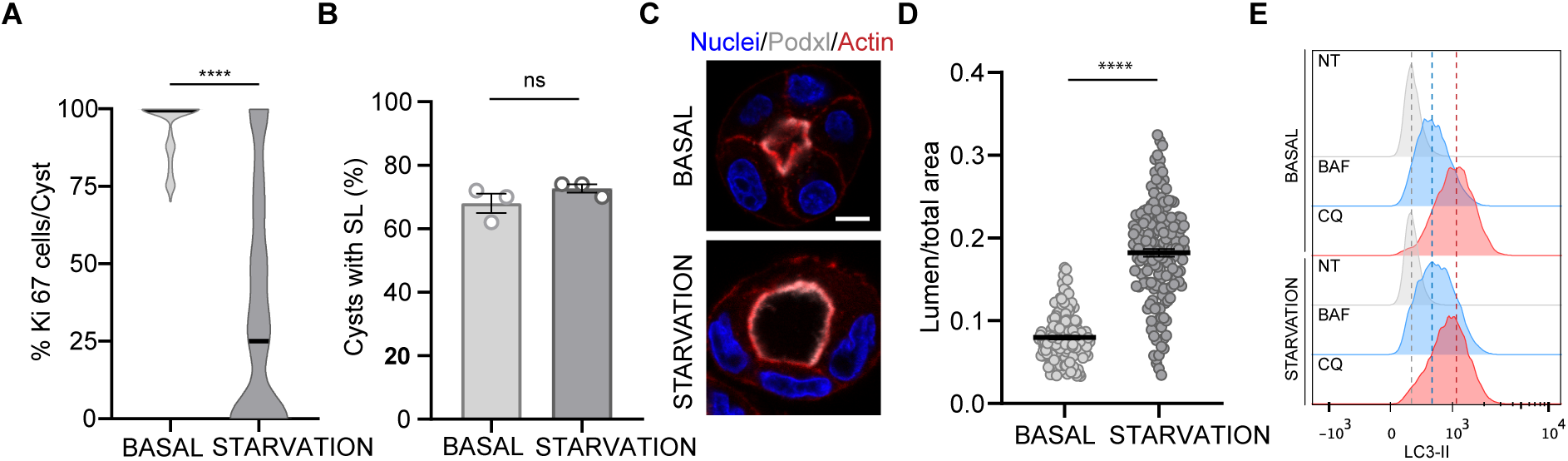
Nutrient deprivation halted MDCK cyst development, leading to the formation of a single large lumen. **(A)** Quantitative analysis of Ki67 in 72h cysts shows a marked decrease in proliferation under starvation conditions (HBSS medium). (N=283 cells). Mann-Whitney test was conducted. **(B)** Percentage (%) of SL in 72h cysts under normal (MEM medium) and starvation (HBSS medium) conditions for 48h. Data are expressed as the mean ± SEM from 3 independent experiments. Unpaired two-tailed t-test was conducted. **(C)** 72h cysts under starvation conditions present enlarged lumens. Scale bar=10 μm. Nuclei are shown in blue. **(D)** Quantitative analysis of the lumen area from IF images shown in panel (C). Data are expressed as the mean ± SEM from 3 independent experiments. (N=330 cysts). Mann-Whitney test was conducted. *P* values= \*\*\*\**p*<0.0001; ns=non-significant.

**FIGURE S6.**
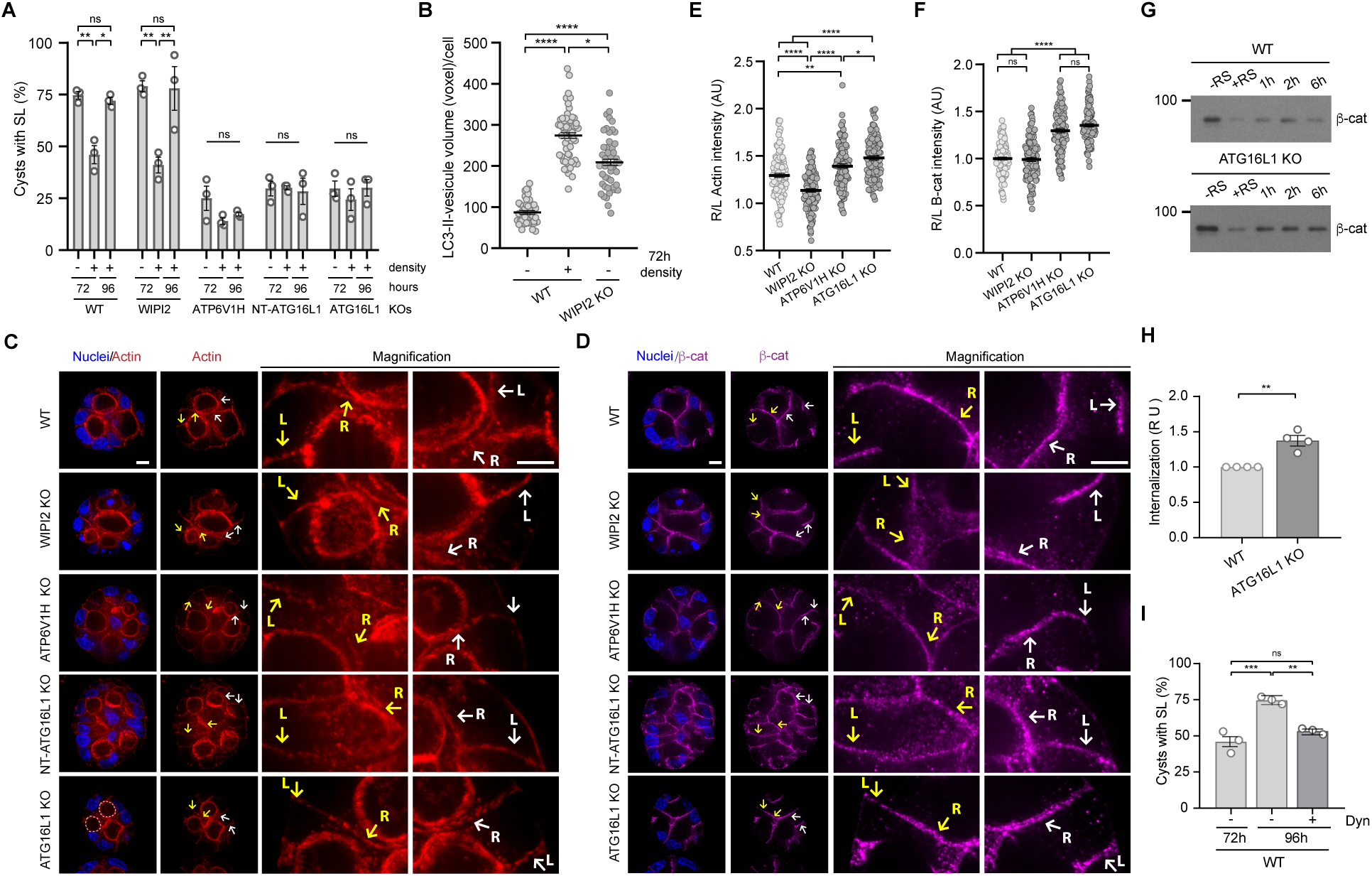
Noncanonical autophagy orchestrates membrane remodeling during MDCK cysts lumen formation. **(A)** Quantitative analysis shows that 96h cysts derived from cells deficient in noncanonical autophagy (ATP6V1H KO, NT-ATG16L1 KO, and ATG16L1 KO), grown at high cell density, fail to resolve multiple lumens. Data are expressed as the mean ± SEM from 3 independent experiments. (N=4500 cysts). One-way ANOVA, with test was performed. **(B)** Quantitative analysis shows that high density WT cyst exhibits an increase in LC3-positive vesicles size, a hallmark of noncanonical autophagy. Data are expressed as the mean ± SEM from 3 independent experiments. One-way ANOVA test was performed. **(C-D)** Actin and β-catenin (β-cat) accumulate at remodeling membranes in 96h cysts lacking noncanonical autophagy (ATP6V1H1, NT-ATG16L1, and ATG16L1 KOs), grown at high cell density. R=remodeling membrane. L=lateral membrane. Nuclei are shown in blue. Scale bars=10 μm and 5 μm (magnification). **(E-F)** Quantitative analysis shows the fluorescence intensity ratio of actin and β-cat between R and L membranes. Data are represented as the mean ± SEM from 3 independent experiments. (N=423 for actin and 446 for β-cat). One-way ANOVA test was performed. **(G)** Biotinylation assays show that noncanonical deficient cells (ATG16L1 KO) retain β-cat in the cytoplasm for longer periods compared to WT, assessed by WB. RS: reducing solution. **(H)** Quantitative analysis from WB shown in panel (B). Data are expressed as the mean ± SEM from 4 independent experiments. Mann-Whitney test was performed. **(I)** Percentage (%) of SL in cysts, grown at high density, shows that cysts fail to form single lumen upon inhibition of endocytic trafficking with Dynasore (Dyn). Data are expressed as the mean ± SEM from 3 independent experiments. One-way ANOVA was performed. *P* values=\*\*\*\**p*<0.0001; \*\*\**p*<0.001; \*\**p*<0.01\**p*<0.05; ns=non-significant.

**FIGURE S7.**
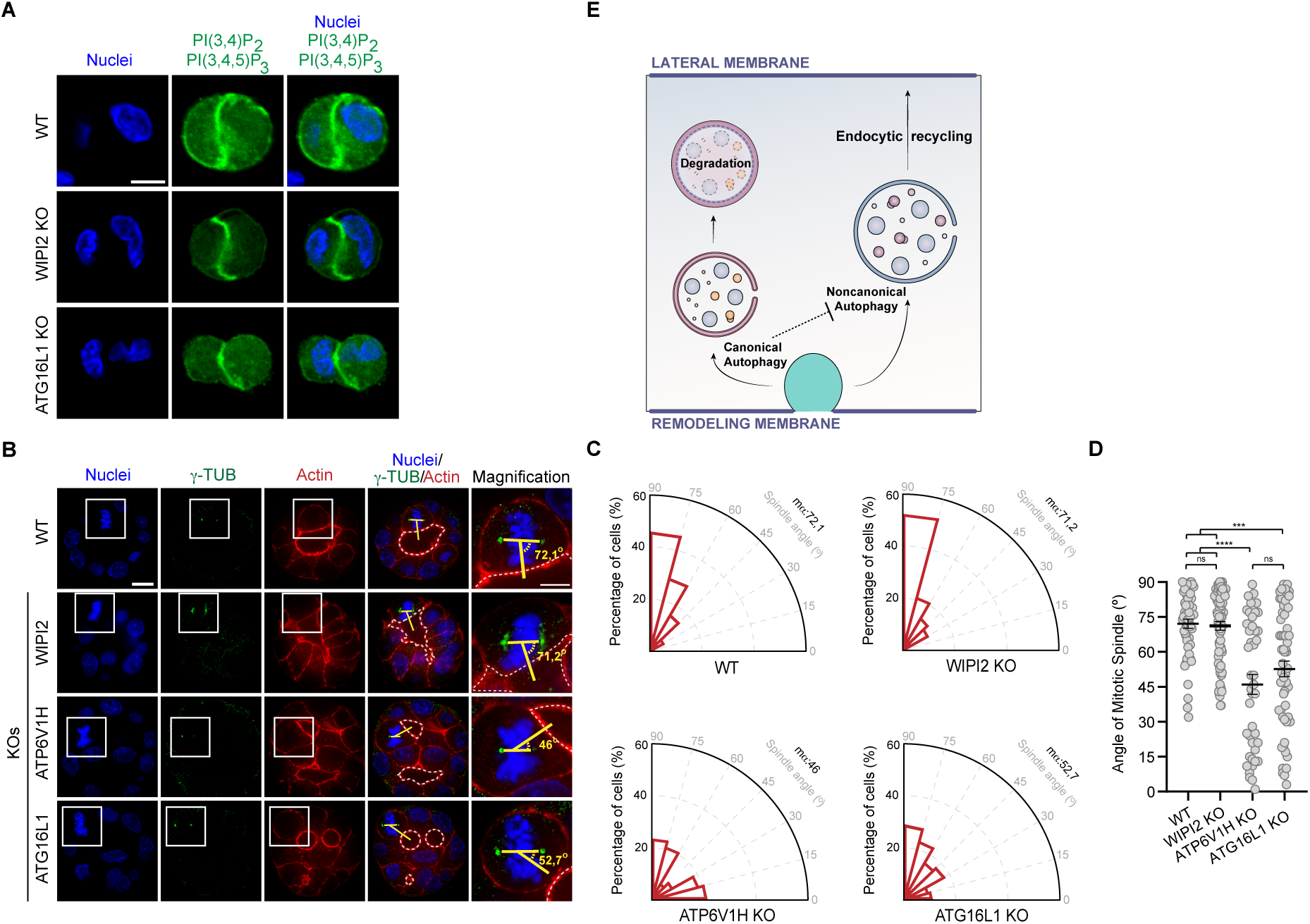
Noncanonical autophagy deficiency leads to spindle defects during MDCK cyst development. **(A)** PI(3,4,5)P3 and PI(3,4)P2 are equally distributed in 24h cyst derived from WT, WIPI2 KO, and ATG16L1 KO cells. **(B)** 72h cysts derived from cells deficient in noncanonical autophagy (ATP6V1H1 KO and ATG16L1 KO) display defects in the angle of cell division. Nuclei are shown in blue. Dotted lines indicate the lumens. Angle of division is shown in yellow. Scale bar=10 μm. **(C)** Radial histogram analysis of division angles analyzed from IF images shown in panel (A). **(D)** Quantification analysis of division angles obtained from IF images shown in panel (A). Data are expressed as the mean ± SEM from 3 independent experiments. Each dot represents one cell (N=282 angles). Kruskal-Wallis test was performed. *P* values= \*\*\*\**p*<0.0001; \*\*\**p*<0.001, and ns=non-significant. **(E)** Schematic model illustrating the contribution of canonical (degradative) and noncanonical autophagy (endocytic recycling of proteins from remodeling membranes to lateral membranes) during lumen resolution in epithelial morphogenesis.

## METHOD DETAILS

### Reagents and antibodies

General laboratory chemicals and reagents were obtained from Sigma-Aldrich and Thermo Fisher Scientific unless specifically indicated. The following reagents were used in this study: Chloroquine (Cytek Biosciences, FCCH100171,) , bafilomycin A1 (Sigma Aldrich, B1793), Phosphatase Inhibitor Cocktail C (Santa Cruz Biotechnology C, 45065), Protease Inhibitor Cocktail (Santacruz, sc-29130), Saponin (Merck, 558255) , Azide (Merck, S2002) .Commercially available polyclonal antibodies (pAbs) or mAbs were used for ATG16L1 (rabbit mAb; Cell Signaling Technology, 8089, and rabbit pAb; MBL life Science, PM040), LC3 (mouse mAb; Cell Signaling Technology, 83506, and rabbit mAb; 3868), WIPI2 (mouse mAb; Abcam; ab105459), ATP6V1H (rabbit pAb; Proteintech, 26683-1-AP), ATP6V1E1 (rabbit pAb, Proteintech, 15280-1-AP), Podocalyxin (rabbit pAb; Proteintech, 18150-1-AP), Claudin-2 (rabbit pAb; Invitrogen, 61-6100), p62 (mouse mAb; Abnova, H00008878 M01), Rab11 (mouse mAb; BD Bioscience, 610656), Sox2 (mouse mAb; Santa Cruz Biotechnology, sc-365823), Oct3/4 (mouse mAb; Santa Cruz Biotechnology, sc-5279), Nanog (mouse mAb; Santa Cruz Biotechnology, sc-293121), CDX2 (mouse mAb; Santa Cruz Biotechnology, sc-60243-1), βCat (rabbit pAb; Santa Cruz Biotechnology, sc-7199), GAPDH (mouse mAb; Santa Cruz Biotechnology, sc-32233), Vinculin (mouse mAb; Sigma Aldrich, v9131), Tubulin (mouse mAb, Sigma Aldrich, T9026), Ki-67 (rabbit mAb; Thermo Fisher Scientific, MA5-14520), Rab5 (mouse mAb; Santa Cruz Biotechnology, sc-46692), Rab7 (mouse mAb, Santa Cruz Biotechnology, sc-376362). Phalloidin labeling probes, and Alexa Fluor 647; were obtained from Sigma Aldrich (TRITC; P1951) and Thermo Fisher Scientific (A22287). Secondary donkey antibodies against mouse or rabbit and labeled with Alexa Fluor 488 (A21202, A21206), Alexa Fluor 555 (A31570), Alexa Fluor 647 (A31573, A31571), or HRP-coupled secondary antibodies (Peroxidase AffiniPure® Goat Anti-Mouse IgG (H+L), 115-035-003, and Peroxidase AffiniPure® Goat Anti-Rabbit IgG (H+L), 111-035-144) were purchased from Thermo Fisher Scientific and Jackson ImmunoResearch Laboratories, respectively.

### Molecular Biology

All DNA primers were obtained from Integrated DNA technologies. pSpCas9(BB)-2A-GFP (PX458) was obtained from Addgene (48138). The following sgRNAs were cloned into the pSpCas9(BB)-2A-GFP (PX458) to generate Knock Outs (KO) in canis familiaris (cf) MDCK cells: cfATG16L1 (5’-ATTCTCTGCATTTAAGCGAT-3’), cfWIPI2 (5’GCTCTTCGCCAACTTCAACC-3’), and cfATP6V1H (5’-GATGACCAAAATGGATATCCG-3’). The sgRNA to generate hATG16L1 KO (5’-ACTGAATTACACAAGAAACG-3’) in hESC was cloned into the pKLV2-U6gRNA5(BbsI)-PGKpuro2ABFP-W (gift from Dr. Bartolomé-Herranz, Instituto de Investigaciones Biomédicas “Sols-Morreale, Spain). The CRISPR Design Tool CRISPOR was used to design the sgRNAs, pMD2.G and pMD-MLVogp plasmids and the DNA constructs expressing Ha or AU1-tagged human ATG16L1 FL (1-607), NT (1-299), and CT (300-607) cloned into retroviral derivative called P12-MMP were king gifts from Dr. Pimentel-Muiños (Centro de Biología Molecular, Spain). The ptfLC3 plasmid was obtained from Addgene (21074). GFP tagged PHD domain construct was used to visualize PtdIns(4,5)P_2_ (Gift from David M. Bryant, University of Glasgow, The United Kingdom).

### Cell culture procedures

MDCK type 2 (MDCK-II) cells were cultured in minimum essential medium (MEM) supplemented with 5% bovine calf serum, 2mM of L-glutamine, and 100U/ml penicillin and 100 mg/ml streptomycin. HEK293T cells were cultured in Dulbecco’s modified Eagle’s medium (DMEM) containing 10% bovine calf serum, 2mM L-glutamine and 100U/ml penicillin and 100 mg/ml streptomycin. H1 hESCs were maintained under feeder-free conditions and passaged on Matrigel-coated plates in mTeSR1 medium. All cell lines were cultured at 37°C in a humidified atmosphere containing 5%CO_2_.

### CRISPR-Cas9 knock out of human and canis familiaris genes and generation of stable cell lines

ATG16L1 KO was generated in a H1 hESC cell line with doxycycline inducible Cas9 expression (Gift from Dr. Bartolomé-Herranz, Instituto de Investigaciones Biomédicas “Sols-Morreale) Spain. hESCs were transfected with the pKLV2-U6gRNA5(BbsI)-PGKpuro2ABFP-W-sgRNA *h*ATG16L1 using Lipofectamine 2000 according to the manufacturer’s protocol. Cas9 expression was induced with doxycycline (2ug/ml) 6 hours before transfection and maintained in the media for 48 hours. ATG16L1 KO, WIPI2 KO, and ATP6V1H KO were generated in MDCK cells. MDCK cells were transfected using Lipofectamine 2000 with SpCas9(BB)-2A-GFP (PX458) plasmids encoding sgRNA targeting cfATG16L1, cfWIPI2, or cfATP6V1H. 48 hours after transfection, single-cell sorting of H1 hESC and MDCK cells was performed on a BD FACsAria cell sorter by gating for BFP*-*positive cells. Individual clones were isolated in 96-well plates and expanded. KOs were confirmed via western blot analysis.

To generate MDCK cell lines stably expressing HA-ATG16L1 FL, HA-ATG16L1 NT, and HA-ATG16L1 CT, ATG16L1 KO cells were infected with HA-ATG16L1 FL, HA-ATG16L1 NT, and HA-ATG16L1 CT retrovirus. Retroviral transductions were performed to express transgenes cloned into the P12-MMP retroviral vector. Virus-containing supernatants were generated by co-transfecting HEK293T cells with the relevant P12-MMP constructs together with helper plasmids expressing gag-pol and env. Infections were carried out by diluting the viral supernatants with fresh medium (1:1) in the presence of polybrene (8 µg/ml), and spinning the resulting mix onto the target cells for 1 h at 2000 rpm, 32 °C. Cells were selected in puromycin (0.5 μg/ml) for two weeks and lysed to evaluate protein expression.

To generate MDCK cell lines stably expressing ptfLC3 or the PI biosensors (GFP-PHD: PI(4.5)P2, AKT-dsRed: PI(3,4,5)P3 and PI(3,4)P2, mNG-PH-SidC:(PI4P)) WT and WIPI2 KO cells were transfected with the ptfLC3 plasmid, and WT, WIPI2 KO, and ATG16 KO cells were transfected with the PI constructs using Lipofectamine 2000. 48 hours after transfection, single-cell sorting was performed on a BD FACsAria cell sorter by gating *GFP or Red-*positive cells. Individual clones were isolated in 96-well plates and cultured in medium supplemented with either Neomycin (500 ug/ml) for ptfLC3-transfected WT and WIPI2 KO cells, or Blasticidin (10 ug/ml) for WT, WIPI2 KO, and ATG16L1 KO cells transfected with the PI constructs. Clones were maintained under selection for two weeks, expanded, and analyzed via western blotting.

### Genetic analysis

Genomic DNA was extracted using a gDNA extraction kit from NZYtech. qPCR analysis was performed to detect the majority of karyotypic abnormalities reported in human ES cells following the manufacturer’s instructions for the hPSC Genetic Analysis Kit (STEMCELL, 07550). Data from AT16L1 KO hESCs were compared to an untargeted H1 hESCs and a genomic sample previously validated as diploid for the analyzed regions using the application available at www.stemcell.com/geneticanalysisapp.

### Pluripotency analysis

For the verification of stem cell pluripotency, the H/M Pluripotent Stem Cell Multi-Color Flow Cytometry Kit (Biotechne, FMC001) was used.

### Western blot

Immunoblots shown are representative of experiments that were repeated and reproduced at least three independent times. Pellets of cells were lysed in lysis buffer supplemented with phosphatase and protease inhibitors (50 mM Tris-HCl pH 7.5, 150 mM NaCl, 0,7% NP40, 0.005% sodium deoxicolate, 0.125% SDS, protease inhibitor cocktail, and phosphatase inhibitor cocktail (Santacruz, sc-29130)) on ice for 30 minutes. Soluble fraction was collected and protein concentrations were measured by Bradford assay using Protein Assay Dye Reagent Concentrate (Bio-Rad, 5000006EDU). For Western blot, proteins were boiled at 95°C for 5 minutes before being loaded onto polyacrylamide gels for SDS-PAGE, separated by SDS-PAGE and transferred to PVDF (Millipore-Sigma, IPFL00010) using a Trans-blot Turbo system (Bio-Rad, 1704150). Membranes were blocked with 5% non-fat dry milk in PBS containing 0.05% Tween-20. All antibodies were diluted in the blocking buffer and incubated at 4 °C on a plate rocker.

### Immunofluorescence

For immunofluorescence of cells in 2d monolayers (over borosilicate coverslips), cysts (ibidi chambers, 80826), neural tube organoids (ibidi chambers, 80826) and micropatterns (cytooplates™ 96 rw cw-s-fn), cells were immunolabeled using protocols adapted from previous studies^52^. Briefly, cultures were fixed in 4% paraformaldehyde (PFA, affimetrix, j19943.k2) for 5–15 minutes at room temperature (rt) and then washed twice in pbs. Samples were blocked for 1 hour in PBS +3% BSA, followed by overnight incubation at 4 °c with primary antibodies diluted in pbs +3% BSA on a gentle rocker. After three washes with PBS, the samples were incubated for 1 hour at Rt with secondary antibodies diluted in PBS and then washed twice in PBS. For anti-LC3 immunolabeling, the protocol was modified. Cultures were fixed with ice-cold 100% methanol for 20 minutes at –20 °c, and both primary and secondary antibody incubations were performed in PBS+3% BSA and 0.1% saponin. Cells cultivated as cysts, were conserved with PBS+0.05% azide at 4 °c for a maximum period of 2 weeks.

Immunostaining of organoid droplets was carried out by first removing the media and rinsing the droplets twice with PBS. The droplets were then fixed in 4% PFA (Sigma, 30525-89-4) prepared in PBS for 1 hour at room temperature. Following fixation, they were washed three times with PBS and permeabilized in PBS containing 0.1% SDS (Merck, 151-21-141) for 1–3 hours, with three additional PBS rinses afterward. Next, the droplets were incubated overnight at 4 °C in a blocking solution composed of 10% BSA (Merck, 9048-144-46-8) in PBT. Using a metal spatula, the droplets were gently detached from the bottom of the 24-well plate and transferred to Eppendorf tubes (Cultek, F7TUBE1EP5-S). They were then incubated with primary antibodies diluted in an antibody solution (1% BSA in PBT) overnight at 4 °C with rotation. After a series of three 5-minute washes with PBS, the droplets were incubated overnight at 4 °C with secondary antibodies in the same solution. Finally, following three additional 10-minute PBS washes, each droplet was transferred to a chamber fitted with a Pwell sticker (iSpacer 0.2 mm, SUNJin Lab, IS016), excess liquid was removed, RapiClear 1.52 (SUNJin Lab, RC147001) was added, and a glass coverslip was placed on top. Zeiss LSM710 and LSM800, Olympus SpinSR, and Nikon AR1+ laser scanning confocal microscopes were used for imaging. Typically, a 40×/0.95 oil-Plan Apochromat or a 60×/1.4 oil-Plan Apochromat objective was employed. Image analysis and composition were performed using ImageJ, FIJI, Imaris, and Huygens.

### LC3 detection and analysis

To detect LC3 (I and II) levels by western blot in hESCs and MDCK cysts, 100 μM chloroquine was added to the media the last 2 hours of incubation at 37°C. After the incubation time, cells were harvested, lysed in lysis buffer, and processed for western blot. To detect LC3-II by IF in hESCs and MDCK cysts, 100 μM chloroquine or 100nm bafilomycin A1 was added to the media the last 2 hours of incubation at 37°C. After the incubation time, cells were fixed and processed for IF. To detect and quantify the ptfLC3 biosensor by flow cytometry, 24hour cysts of WT and WIPI2 KO MDCK cells stably expressing ptfLC3 were harvested, and the MFI of GFP and RFP was measured. Data were acquired using a Cytek Aurora full-spectrum flow cytometer and analyzed with FlowJo software.

To detect LC3-II by flow cytometry in MDCK cysts, 100 μM chloroquine or 100nm bafilomycin A1 was added to the media the last 2 hours of incubation at 37°C when required. Cells were then labelled with LIVE/DEAD® Fixable Near-IR Dead Cell Stain Kit 633-635 nm (ThermoFisher, L10119) for 10 minutes to separate dead cells for the analysis. After the incubation time, endogenous intracellular vesicular LC3 levels (LC3-II) were detected using the Guava Autophagy LC3 antibody-based assay Kit (Luminex, FCCH100171). Manufacture instructions were followed. The quantification of the relative contributions of canonical versus noncanonical autophagy during MDCK cysts development was achieved by establishing the difference between the mean fluorescence intensity of LC3 (MFI LC3) with either chloroquine (CQ) or bafilomycin A1 (BAF) and MFI LC3 before treatment (NT); normalized to MFI LC3 before treatment (NT). In this case we obtained a value for the total autophagy flux [TA flux=(MFI LC3CQ-MFI LC3NT)/MFI LC3 NT]; canonical autophagy flux [CA flux=(MFI LC3BAF-MFI LC3NT)/MFI LC3 NT]; or the noncanonical autophagy flux as a result of their subtraction (NCA flux= TA-CA). Data were acquired on FACSCanto II Flow Cytometer analyzed with FlowJo software.

### Generation of spheroids and micropatterns

To cultured MDCK-II as 3D cysts, single cell suspension (2 x 10^3^ cells ml^-1^ for low density and 7.5 x 10^4^ cells ml^-1^ for high density) in medium contain 2% Matrigel (BD biosciences, 354234) were plated into μslide 8-well coverglass chambers (IBIDI, 80827,), pre-coated with 5ul of 100% Matrigel and allowed to solidify at room temperature for 30 minutes. Cysts were grown using normal MDCK-II culture conditions and fixed at indicated time points.

To prepare MDCK-II tubes in micropatterned-platforms, extracellular matrix-coated surfaces were placed in a 6-well plate and rinsed with PBS and incubated with culture medium for 1 hour at 37°C. Cells were trypsinized to a single cell suspension of 6 × 10^4^ cells, seeded in their appropriate culture medium and incubated for 1 hour at room temperature and 2 hours at 37°C in a humidified atmosphere containing 5% CO_2_. Once cells are completely attached, the culture medium is replaced with Matrigel-containing medium. MDCK-II medium is supplemented with 3% FBS and 3% Matrigel. Chips are kept at 37°C in a humidified atmosphere containing 5% CO_2_. The medium was changed 2 days and grown for 4 days until tubes with lumen formed.

### Secondary neurulation

To study epithelialization and lumen resolution in neural tube organoids, solid aggregates were generated before Matrigel induction using a two-day spheroid preassembly protocol, following previously published methodology^7^. First, cells were rinsed twice in PBS and incubated in 0.5 mM EDTA at 37°C for 4 minutes to facilitate detachment, after which the resulting multicellular clumps were collected in mTeSR1. Approximately 10⁶ cells forming clumps were then seeded into a six-well plate with mTeSR1 and maintained under shaking conditions at 300 rpm, 37°C, and 5% CO₂. The next day, 3 µM CHIR99021 (Millipore, 361571) was added to the medium and left another 24 hours under shaking conditions. Subsequently, spheroids from each well were collected, centrifuged, and resuspended in 50–80 µL of Matrigel (Corning, 356230) to generate droplets. A 10 µL droplet of the Matrigel-cell suspension was pipetted into each well of a twenty-four-well tissue culture plate (Corning, 353046) and incubated for 13 minutes at 37°C. Next, an N2B27-based neural induction medium was added, consisting of a 1:1 mixture of Advanced DMEM/F12 (Gibco, 31331-028) and Neurobasal medium (Gibco, 21103-049), supplemented with 0.5× N2 (Gibco, 17502-001), 0.5× B27 (Gibco, 17504-001), 1× nonessential amino acids (Gibco, 1140-035), 1× sodium pyruvate (Gibco, 11360-088), 0.5× GlutaMax (Gibco, 35050-038), and 0.1 mM β-mercaptoethanol (Gibco, 31350-010). To induce posterior neuromesodermal differentiation, the medium was further supplemented with 3 µM CHIron, 0.1 µM LDN193189 hydrochloride, 0.1 µM All-Trans Retinoic Acid, and Human FGF-basic (FGF-2/bFGF) (aa 10-155) Recombinant Protein (1/1000). The organoids were cultured under these conditions for 48 hours, after which CHIron, LDN193189 hydrochloride, All-Trans Retinoic Acid, and Human FGF-basic were removed from the medium. Culture medium was changed daily, and plates were maintained at 37°C with 5% CO₂.

ES cells were routinely cultured in gelatin-coated plates in Fc medium supplemented with 2i/LIF at 37°C, 5% CO_2_, 21% O_2_. Cells were routinely tested for mycoplasma contamination. For embryoid formation, mES cells (20,000 cells/well) were suspended in a complete medium containing 5% Matrigel (Corning, Life Sciences). Cell suspension was plated in an 8-well culture slide (Corning, Life Science) on a thin layer coating of 100% Matrigel. The plates were kept at 37°C in a humidified atmosphere of 5% CO_2_ for 72h and the medium was changed every 2 days.

### Biotinylation assay

For the MDCK endocytosis assay, cells were seeded in p60 dishes pre-coated with Matrigel and used once they reached 60–70% confluence. Initially, cells were washed three times with cold (4°C) PBS containing Ca²⁺ and Mg²⁺, and then incubated for 20 minutes at 4°C with 1 mg/ml Sulfo-NHS-SS-Biotin (Pierce, PG82077) in the same PBS/Ca/Mg solution. After biotinylation, the dishes were washed three times with 100 mM lysine in PBS/Ca/Mg. Control samples were immediately lysed in the lysis buffer specified for the biotinylation assay and kept on ice. For the remaining samples, pre-warmed culture medium (37°C) supplemented with 20 mM HEPES (pH 7.4) was added, and the cells were incubated at 37°C for the indicated time points. Following the incubation periods, cells were transferred back to 4°C to halt intracellular trafficking and then treated twice (each for 8–20 minutes) with a reducing solution composed of 50 mM glutathione, 75 mM NaCl, 1 mM EDTA (pH 8), 1% BSA, and 75 mM NaOH (adjusted to pH 7.4). This treatment selectively removed the reversible biotin label from surface proteins, ensuring that only internalized proteins remained biotinylated. Finally, cells were incubated with 5 mg/ml iodoacetamide in PBS/Ca/Mg for 5 minutes, washed twice with PBS/Ca/Mg, and then lysed. Biotinylated proteins were isolated via pull-down using neutravidin-agarose (Thermo Scientific, 29204) as described previously, and both lysates and pull-down pellets were analyzed by western blot.

### Single lumen/ multiple lumen quantifications

Neural tube organoids and MDCK spheroids displaying a single actin/Podocalyxin-positive apical surface with β-catenin/actin staining oriented toward the ECM were classified as having a single lumen. Experiments in which fewer than 50% of structures exhibited normal lumens at 72 hours under control conditions—typically due to suboptimal Matrigel gelification—were excluded and repeated. In MDCK micropatterns, structures with either one or two lumens were quantified as having a single lumen, as previously described. Tubes with one or multiple lumens were considered polarized, whereas structures with a 2D morphology or inverted polarity were classified as non-polarized.

### Imaging and Quantification

Low-magnification fields were randomly selected for initial imaging, followed by higher magnification imaging for detailed quantification. Immunofluorescence experiments were performed in at least three independent replicates, and representative images were selected from these samples for analysis. Prior to quantification, background subtraction was carried out in Fiji. Projections of individual cells were generated, and fluorescence intensity was quantified as either integrated density per cell or total cyst volume. Actin and β-catenin signal polarization was assessed by measuring the ratio of the resolving membrane signal (i.e., the signal of actin or β-catenin between two un-fused lumens) to the lateral membrane (actin or β-catenin signal) of the same cell. Circularity was determined using Fiji’s “Analyze Particles” tool, calculated as 4π(Area)/(Perimeter²), with values closer to 1 indicating a shape that is nearly perfect. Lumen space percentage was computed by manually outlining the total spheroid area with the Freehand Selection tool and calculating the ratio of lumen area to total spheroid area. For ATG16L1 KO spheroids, lumen space percentage and volume were alternatively measured using CartoCell^42^, a deep-learning-based image analysis pipeline. All images were acquired under identical settings, including laser power, gain, and exposure time.

### Intensity profiles

Fluorescence intensity profiles were generated using ImageJ by drawing a line across the region of interest and measuring the pixel intensity along the line.

### Vesicle Quantification

For vesicle number and intensity quantification, raw image stacks were initially processed using Huygens Professional software. Deconvolution was performed with the Classic Maximum Likelihood Estimation (CMLE) algorithm using a theoretical point spread function (PSF) to enhance resolution and signal clarity. Vesicle detection and segmentation were achieved using the Object Analyzer tool in Huygens. Vesicles positive for LC3, Rab11 or other markers were identified based on intensity, size, and shape parameters. Intensity thresholding distinguished vesicles from background fluorescence, and size filtering excluded artifacts below a predefined minimum volume threshold. Shape analysis further ensured the accurate identification of vesicular structures. For each region of interest (ROI) or individual cell, vesicle number was determined as the total count per ROI, while vesicle volume and fluorescence intensity were assessed based on mean and total values per vesicle.

### Measurement of Spindle Orientation

Images of mitotic cells stained with acetylated α-tubulin mAb were collected. A line was drawn connecting the two spindle poles to define the spindle axis, and a second line was drawn from the center of the apical membrane to the midpoint of the basal membrane, delineating the apicobasal axis. The angle between these two lines was then measured to determine spindle orientation.

### An Agent Base model to study balance between canonical and noncanonical autophagy

These types of models, composed of agents that behave following certain rules defined to simulate the behavior of biological entities, are commonly used as a framework to test hypothesis and define the minimal sets of rules that can reproduce a biological response of interest. In this case, the model is designed to simulate a single cell where two locations are established, representing two regions of the extracellular membrane. Autophagy is simulated by defining a number of endocytic vesicles (the agents in our model) that are generated at one of the membranes and follow the rules defined programmatically, inspired by the experimental observations. Since both canonical and noncanonical vesicles share many components, it is expected some inter-dependence between the fluxes of canonical and noncanonical autophagy. The model is designed to be used to test these potential dependencies, and to make predictions about how variation in one flux affect the other. The basic scheme of the model is the following: (1) Endocytic vesicles (initially colored as green) are generated in one of the membranes and enter the cell. (2) The vesicles acquire an identity as canonical (blue) or noncanonical (noncanonical), based on rates defined by the user. (3) Canonical vesicles are defined as following the expected canonical autophagy path and are degraded after a given time inside the cell; (4) noncanonical vesicles on the other hand, follow the noncanonical autophagy path, assumed as a transport mechanism of certain cargo to another location of the cell (in this case the other location in the membrane). (5) The number of vesicles of each type at each time point is quantified and used to compare with the experimental fluxes measured experimentally. Code used to run the simulations is included as supplementary information in the form of a Jupyter notebook running using Julia programming language. https://nextjournal.com/a/RQL9FiFzKN4ZdCbzcQK4dE?token=Txke4bC58AoazKm9zqZ7QA

### Statistical analysis

Statistical analysis was performed with GraphPad Prism (version 8). Data are expressed as mean ± S.E.M. (standard error of the mean). Outliers were excluded by the ROUT method (5%). Comparisons for two groups were calculated using unpaired two-tailed Student’s *t*-tests (for two groups meeting the normal distribution criteria) or Mann-Whitney U test (for two groups without normal distribution) according to the Shapiro–Wilk normality test. Comparisons for more than three groups were calculated using One-way ANOVA (for three or more groups meeting the normal distribution criteria) or Kruskal-Wallis test (for three or more groups without normal distribution). Statistically significant differences are denoted on the figures as: ∗*p*<0.05, ∗∗ *p*<0.01, ∗∗∗ *p*<0.001, ∗∗∗∗ *p*<0.0001.

## Notes

### Competing Interest Statement

The authors have declared no competing interest.

